# A screen to identify antifungal antagonists reveals a variety of pharmacotherapies induce echinocandin tolerance in *Candida albicans*

**DOI:** 10.1101/2025.02.18.638903

**Authors:** Parker Reitler, Christian A. DeJarnette, Ravinder Kumar, Katie M. Tucker, Tracy L. Peters, Nathaniel R Twarog, Anang A. Shelat, Glen E. Palmer

**Affiliations:** Department of Clinical Pharmacy and Translational Science, College of Pharmacy, University of Tennessee Health Sciences Center, Memphis, USA; Department of Pharmacy and Pharmaceutical Sciences, St Jude Children’s Research Hospital, Memphis, TN, USA; Department of Chemical Biology and Therapeutics, St Jude Children’s Research Hospital, Memphis, TN, USA

## Abstract

Through screening a comprehensive collection of drugs approved for human use, we identified over 20 that oppose the antifungal activity of the echinocandins upon the infectious yeast, *Candida albicans*. More detailed evaluation of five such drugs, including the atypical antipsychotic aripiprazole and the tyrosine kinase inhibitor ponatinib, indicated they promote *C. albicans* survival following exposure to the echinocandin antifungals. The activity of the five selected antagonists was dependent upon the Mkc1p MAPK pathway, however, ponatinib was paradoxically shown to suppress phosphorylation and therefore activation of Mkc1p itself. Components of several other signaling pathways are also required, including those of calcineurin and casein kinase-2, suggesting the observed antagonism required much of the cell wall stress responses previously described for *C. albicans*. Transcriptome analysis revealed that the antagonists stimulated the expression of genes involved in xenobiotic and antifungal resistance, and suppressed the expression of genes associated with hyphal growth. Thus, the echinocandin antagonistic drugs modulate *C. albicans* physiology in ways that could impact its pathogenicity and/or response to therapeutic intervention. Finally, a mutant lacking the Efg1p transcription factor, which has a central role in the activation of *C. albicans* hyphal growth was found to have intrinsically high levels of echinocandin tolerance, suggesting a link between modulation of morphogenesis related signaling and echinocandin tolerance.

**Importance:** We report a substantial number of previously unknown drug interactions that modulate the echinocandin sensitivity of one of the most prevalent human fungal pathogens, *Candida albicans*. The echinocandins are the first line therapy for treating disseminated and often lethal *Candida* infections, that account for >75% of invasive fungal infections in the U.S.. For largely unknown reasons, a substantial number of patients with invasive candidiasis fail to respond to treatment with these drugs. The finding of this study suggest that co-administered medications have the potential to influence the therapeutic outcomes of invasive fungal infections through modulating antifungal drug tolerance and/or fungal pathogenicity. The potential for echinocandin antagonistic medications to influence therapeutic outcomes is discussed.

## Introduction

The CDC estimates the direct healthcare costs of treating fungal disease to be over $7 billion each year in the U.S. alone [1]. Infections involving fungal invasion to the deeper organs are of greatest concern, and predominantly occur in immunocompromised individuals. Several *Candida* species account for more than 75% of disseminated fungal infections in the U.S. [2, 3] with attributable mortality rates of 35-75% [4–6]. They are also a common cause of mucosal diseases including oral and vaginal thrush with *Candida albicans* the most prevalent and virulent [7]. The echinocandins are the first line therapy for invasive *Candida* infections, and disrupt fungal cell wall synthesis through inhibiting β (1,3)-D-glucan synthase [8]. However, nearly a third of patients with invasive candidiasis (IC) fail to respond to treatment with an echinocandin [9–12], and outcomes are significantly worse for neutropenic patients [13]. The incidence of echinocandin resistance is < 0.5% for *C. albicans* [14, 15], thus, genetically encoded antifungal resistance does not account for the majority of treatment failures [16–19]. Several host-related factors have been proposed to account for the discordance between *in vitro* susceptibility tests and patient outcomes [18]. Certainly, rapid diagnosis and administration of an appropriate antifungal agent can reduce patient mortality [20, 21]. The severity of patient immunosuppression may also impact the response to antifungal therapy [19, 22]. Either way, many treatment failures remain unexplained. Given the limited therapeutic options available, it is critical to identify and address the causes of therapeutic failure to improve patient outcomes.

As natural inhabitants of the human gastrointestinal and reproductive tracts, several medically important *Candida* species are routinely exposed to medications and other pharmacologically active substances consumed by their human host. Given the fundamental conservation of metabolic and signaling modules between eukaryotic genera, it is likely that many drugs designed to modulate human physiology may dysregulate analogous pathways in resident or infecting fungi [23–26]. Yet, the influence of most medications upon fungal physiology, pathogenicity and antifungal susceptibility, remains largely uncharacterized. Moreover, it is unclear if such drug-fungus interactions have the potential to influence the incidence or outcomes of mycotic disease. We and others have previously reported that a wide variety of drugs approved for human use can diminish the capacity of the azole antifungals to inhibit the growth of pathogenic *Candida* species [27, 28]. We also identified a smaller collection that moderate *C. albicans* sensitivity to the echinocandin, caspofungin [27]. These interactions are potentially important determinants of clinical outcome, as chemically induced antifungal resistance occurring within a patient receiving an antagonistic medication would not be detected by *in vitro* antifungal susceptibility testing methods. However, the drug collections screened in previous studies only offered partial coverage of the current pharmacopeia. Additionally, the physiological impact of these drug-fungus interactions and their potential to impact interaction with the mammalian host remains unknown. The purpose of this study was to identify medications that alter *C. albicans* sensitivity to the echinocandins and examine the mechanisms by which they act.

## Materials and Methods

### Growth conditions

*C. albicans* was grown on yeast extract-peptone-dextrose (YPD) agar plates at 30°C and incubated in YPD liquid medium unless otherwise stated. Select strains were graciously provided either by Dr. Brian M. Peters or Dr. Todd B. Reynolds. Kinase defective mutants were kindly provided by Dr. Damian Krystan [29].

### Drug stocks

Stock solutions of each antifungal (caspofungin, micafungin, and anidulafungin), as well as each compound (aripiprazole, cinacalcet, haloperidol, netupitant, and ponatinib) were prepared at 10 mM in dimethyl sulfoxide (DMSO) and diluted to required working concentrations. The FDA collection was purchased from SelleckChem with all 2,701 compounds dissolved in respective solvent listed (either DMSO or water) at 10 mM in 384-well plates. Each was further diluted to 200 µM, and 1-μl volumes were dispensed into the flat-bottomed 384-well polystyrene plates that were used for the chemical screens.

### Antifungal antagonism screens

Assays were performed similarly as described [28]. The wells of the 384-well flat-bottom assay plates were seeded with 1 μl of the 1 mM stock solutions of each library compound in DMSO or with DMSO alone (control). SC5314 was grown overnight in YPD medium at 30°C, and the cells were harvested by centrifugation, washed twice in phosphate-buffered saline (PBS), and resuspended at ∼6.25 × 10^4^ cells/ml in Roswell Park Memorial Institute (RPMI) 1640 medium buffered to pH 7 with (3-(N-morpholino)propanesulfonic acid) (MOPS) 2% glucose supplemented with the indicated concentration of caspofungin. A 39-μl volume of the cell-plus-antifungal drug suspension was then dispensed into each well of the drugged 384-well plates to achieve approximately 1000 cells/well. Control wells had antifungal drug but no test/library compound (antifungal alone) or had neither the antifungal nor the library compound (minus drug/untreated control). The final concentration of DMSO was 0.55% in all wells, with each library compound supplied at a final concentration of 5 μM. After 72 hours of incubation at 35°C, growth was measured as OD_600nm_ without shaking. Wells with visible precipitation were excluded from analysis. Growth was compared to the antifungal alone and untreated control and expressed as percent growth. The growth of each drug-treated well was also compared to antifungal alone wells, and the Z-score was calculated [30]. “Hits” were identified as compounds with a Z-score of ≥ 3 in two independently run screens.

### Antifungal susceptibility assays

Antifungal susceptibility testing was performed using the broth microdilution method as described in CLSI document M27-A3 [16], with minor modifications. Each echinocandin was diluted in DMSO, resuspended in RPMI-pH 7 (2% glucose) at twice the final concentration, and serially diluted. *C. albicans* strains were grown overnight in YPD at 30°C, resuspended at 1 x 10^4^ cells/ml in RPMI-pH 7, and 100 µl transferred to wells of a round bottom 96-well plate containing and equal volume of diluted echinocandin solution. The final concentration of DMSO was 0.5% for all treatments, with drug-free control wells having DMSO alone. Plates were incubated without shaking for 72 hours at 35°C, with plates scanned every 24-hours using an EPSON Perfection v700 Photo scanner. Experiments were performed in biological duplicate.

### Growth kinetic assays

Growth curve assays were set up in 96-well plates using RPMI-pH 7 (2% glucose). *C. albicans* strains were grown overnight in YPD at 30°C and the cell density was adjusted to 1 x 10^4^ cells/ml in the appropriate medium for the growth kinetic assays. An aliquot of cell suspension (100 µl) with an equal volume of twice the desired drug concentration of each antagonist. Cells were then incubated at 35°C inside a BioTek Cytation 5 plate reader shaking for 48 hours, and OD_600nm_ read every 30 min. Data was then analyzed via GraphPad Prism software. These assays were repeated in biological duplicate.

### Cell survival and cell damage assays

For cell survival assays, the C. albicans parental strain GP1 and strain CAI4+pKE1-NLUC were grown overnight in YPD at 30°C. Cells were washed and subcultured at 1 x 10^6^ cells/ml in 10 mls of RPMI-pH 7 (2% glucose) supplemented with 0.5% DMSO (vehicle), 0.4 µM or 1.6 caspofungin alone or in the presence of 5 µM aripiprazole, cinacalcet, haloperidol, netupitant, or 2.5 µM ponatinib. Cells were incubated at 35°C for 6 hours plus or minus drug at shaking. After incubation, cells were spun down at 3000 revolutions per minute (rpm) for 5 minutes, and washed twice with sterile dH_2_O. Cells were then resuspended in sterile dH_2_O, serially diluted, and 100 µl of cell solutions were plated onto YPD plates. Plates were then incubated for 48 hours at 30°C, and cell viability was measured by counting colony forming units (CFU). For cell damage assays, cells were grown under the same conditions, and a portion of supernatant was collected for damage Experiments were repeated in biological triplicate.

For cell damage assays, supernatant was collected from the cells grown in the previously listed condition. Luciferase based assays were performed as previously described [31]. Experiments were performed in biological triplicate and statistical significance was calculated via one way Variance of Analysis (ANOVA) test.

### RNA sequencing analysis

SC5314 was grown in YPD medium at 30°C overnight, then sub-cultured at 1 x 10^6^ cells/ml into 50 ml RPMI-pH 7 medium supplemented with either 5 µM aripiprazole, cinacalcet, haloperidol, netupitant, 2.5 µM ponatinib, or with 0.5% DMSO (vehicle control). Cells were then incubated at 35°C for 6 hours with shaking. Cells were then harvested at 4°C, supernatant removed, and cells frozen at −80°C. Total cellular RNA was extracted using the hot phenol method [32]. Novogene provided RNA library preparation and sequencing analysis as fee-for-service. Messenger RNA was purified from total RNA using poly-T oligo attached magnetic beads. After fragmentation, the first strand cDNA was synthesized using random hexamer primers, followed by the second strand cDNA synthesis using either dUTP for directional library or dTTP for non-directional library. Library was checked with Qubit and real-time PCR for quantification and bioanalyzer for size distribution detection. Libraries were then pooled and sequence on Illumina platforms. Genes were mapped to the SC5314 haploid genome assembly 22. Drug responsive genes were identified as those significantly up or downregulated compared to their respective vehicle controls, with significant gene expression identified as either > 1 or < -1 log_2_fold (p-value < 0.05, adjusted p-value < 0.05 to account for false discovery rate). Samples were prepared in independent biological duplicate, and log_2_fold was converted to fold change, averaged, and the average value was converted to log_2_fold for the term “AVG log_2_fold”.

### Western blot analysis

SC5314 strain was inoculated in YPD and incubated overnight at 30°C. The next day, SC5314 was subcultured into 75 mls of pre-warmed RPMI-pH 7 (2 % glucose) supplemented with vehicle, 5 µM of aripiprazole, cinacalcet, haloperidol, netupitant, 2.5 µM ponatinib alone or in combination with caspofungin. SC5314 was washed with PBS and diluted to an OD_600nm_ of 0.25 in RPMI-pH 7 medium using a spectrophotometer (BioPhotometer from Eppendorf). Flasks were incubated at 35°C with 250 rpm shaking. After two hours of growth, cells were harvested at 4000 rpm for 30 minutes and supernatant removed. The cells were resuspended in PBS, transferred into a 1.5 mL screw-capped tube, and harvested at 13000 rpm for 10 minutes. Tubes were immediately stored at -80°C.

### Protein extraction

Cell were removed from -80°C, thawed on ice, and then resuspended in 120 ml buffer containing (300 mM NaCl, 10 mM Tris, pH 8.0, 0.1% NP-40, and 10% glycerol) and containing 1X protease inhibitor cocktail, phosphatase inhibitor cocktail set II (Millipore Sigma, 1:100 dilution), and 1 mM DTT [33]. Glass beads equivalent to the size of a pellet were added into tube and tube was kept in bead beater Bullet Blender Gold (Next Advance) operated for 5 cycles of 1 min each, with 1-minute incubations on ice for each cycle. After cell lysis, whole cell extract was collected in a fresh 1.7 ml Eppendorf tube and kept on ice. Protein quantification was performed by Bradford assay (Thermo Fischer Scientific, cat # 1856210), and protein concentration was calculated comparing the result to a standard curve.

### SDS-PAGE and immunoblotting

30 µg of protein sample was mixed with 4X loading buffer (from Bio-Rad, cat #1610747), and protein samples were heated for 5 minutes at 100°C in a thermocycler. Protein samples were cooled to room temperature and loaded onto a 10 % SDS-PAGE gel. The gel was run at a constant 100 V until the dye front reached the bottom of the gel. Proteins were transferred onto nitrocellulose membrane (0.2 uM from Bio-Rad, cat #162-0112) using wet transfer for 1 hour at constant 100 V. After transfer, membranes were rinsed with distilled water and stained with Ponceaus S stain (from Thermo Fisher Scientific, cat # A40000279) and blots were imaged after rinsing with distilled water. Blots were then incubated in blocking buffer in tris-buffered saline with 0.1% Tween 20 detergent (TBST) pH-7.5 (blocking grade blocker from Bio-Rad cat # 170-6404) (TBST diluted from 10X stock from Bio-Rad cat # 170-6435) for 1 hour at room temperature. Blots were then incubated overnight with rabbit anti-phospho-p44/42 antibody (from Cell Signaling, cat # 4370) diluted to 1:2000 in TBST with (5 % nonfat skimmed milk powder in TBST pH-7.5) at 4°C shaking overnight. The next day, primary antibodies were removed, and blots were washed with TBST for 10 minutes three times. After the final wash, blots were incubated for 1 hour at room temperature with HRP conjugated goat anti-rabbit IgG (H+L) (from Invitrogen cat # 31460) diluted to 1:10000 in TBST with 5% nonfat skimmed milk powder. After incubation, nonbound antibodies were removed at blots were washed thrice as above. Finally, the signal was detected using ECL (SuperSignal West Femto Luminol/Enhancer Solution and SuperSignal West Femto Stable Peroxide Buffer from Thermo Fischer Scientific), and images were captured using Molecular Imager ChemiDoc XRS+ with Image Lab Software (from Bio-Rad). Experiments were performed in biological quadruplicate and statistical significance was calculated via one-way ANOVA test.

## Results

### Several drugs approved for human use oppose the antifungal activity of echinocandins upon *Candida albicans*

To identify medications that potentially modulate the efficacy of echinocandin therapy, we conducted a simple screen. *C. albicans* (strain SC5314) was suspended in RPMI medium supplemented with approximately 4X the MIC (0.78 µM) of caspofungin, and dispensed into 384-well plates arrayed with a collection of 2701 pharmacologically active small molecules - most of which are approved for human use by the FDA (Food and Drug Administration), EMA (European Medicines Agency), and other approval agencies. Each compound was provided to a final concentration of 5 µM, and control wells contained either no drugs or antifungal (DMSO vehicle growth control), or caspofungin alone. Given the cidal mode of action of the echinocandins, we measured fungal growth in each well as OD_600nm_ after 72-hours incubation at 35°C. This strategy was intended to identify drugs that promote fungal survival in the presence of caspofungin (i.e. tolerance), as well as any that induce outright resistance (i.e. insensitivity). Compounds were called as hits if they restored fungal growth in the presence of caspofungin (Z-score ≥ 3) in each of two independent replicate screens. A total of 22 hits, corresponding to 21 distinct chemical entities, were identified as opposing the antifungal activity of caspofungin upon *C. albicans* (table 1). This included: 1. aripiprazole, an atypical antipsychotic that we had previously reported to diminish the antifungal activity of the azole antifungals upon *C. albicans* [34]; 2. cinacalcet, a calcimimetic used to treat hyperparathyroidism; 3. the antipsychotic haloperidol; 4. the antiemetic netupitant; and 5. ponatinib, a multi-targeted tyrosine-kinase inhibitor used to treat leukemia. These 5 drugs were selected as representative hits for further evaluation, and their activity compared.

**Table 1.** List of caspofungin antagonists identified from the SelleckChem FDA collection. Wild-type C. albicans strain SC5314 was dispensed in RPMI-pH 7 2% glucose with the SelleckChem FDA collection. Screens were performed with 4X MIC of caspofungin (CAS) into 384-well plates arrayed with each library compound at a final concentration of 5 μM. Plates were incubated for 48 hours at 35°C and then growth quantified as OD_60onm_-Growth was calculated to untreated control wells (DMSO alone) relative to antifungal alone and defined (% growth). Hits were called as compounds with Z-score of ≥ 3 compared to cells + antifungal alone in two independently run experiments.

| AVG % Growth<br>(untreated<br>control) | AVG Z-Score<br>(CAS alone) | Product Name | Target/Mechanism | Pathway |
| --- | --- | --- | --- | --- |
| 43.914 | 29.886 | Ponatinib (AP24534) | Bcr-Abl:FGFR:PDGFR:VEGFR | Angiogenesis |
| 45.109 | 28.358 | Haloperidol | Dopamine Receptor | Neuronal Signaling |
| 29.431 | 15.090 | Sanguinarine chloride | phosphatase | Others |
| 22.266 | 13.522 | Cinacalcet HCl | CaSR | GPCR & G Protein |
| 3.398 | 13.233 | Oxaliplatin | DNA/RNA Synthesis | DNA Damage |
| 18.920 | 11.505 | Aripiprazole | 5-HT Receptor | Neuronal Signaling |
| 19.965 | 11.413 | Netupitant | Substance P | Others |
| 21.455 | 11.093 | Sanguinarine | Antioxidant | Cancer |
| 13.656 | 9.478 | Fesoterodine Fumarate | AChR | Neuronal Signaling |
| 14.101 | 8.862 | Proanthocyanidins | Others | Others |
| 11.202 | 7.818 | Costunolide | Telomerase | DNA Damage |
| 12.329 | 6.643 | Vorapaxar | Protease-activated Receptor | GPCR & G Protein |
| 9.460 | 6.368 | Bambuterol HCl | Adrenergic Receptor | GPCR & G Protein |
| 10.059 | 6.291 | Methyldopa | 5-HT, Adrenergic, and Dopamine<br>Receptors | Metabolism |
| 10.856 | 6.016 | Acarbose | Others | Others |
| 8.516 | 5.029 | Suprofen | COX | Neuronal Signaling |
| 8.226 | 4.981 | Nicardipine HCl | Calcium Channel | Transmembrane<br>Transporters |
| 7.214 | 4.811 | Biotin (Vitamin B7) | Vitamin | Metabolism |
| 6.322 | 3.852 | Iopamidol | Others | Others |
| 5.740 | 3.583 | Yangonin | Others | Others |
| 5.391 | 3.414 | Clindamycin alcoholate | Anti-infection | Microbiology |
| 5.104 | 3.150 | Bethanechol chloride | AChR | Neuronal Signaling |

Initial dose-response assays confirmed that all 5 drugs opposed the antifungal activity of caspofungin, although the degree of protection conferred by each varied. In all cases, their protective effect upon *C. albicans* was progressively more apparent at later time points (i.e. 48- and 72-hours) (figure 1A). When read after 24-hours of incubation, caspofungin MICs were comparable for all treatments, but residual growth was apparent at concentrations above the MIC in the presence of all 5 antagonists. Additionally, with the exception of ponatinib, the other four antagonists appear to preferentially restore *C. albicans* growth at the highest concentrations of caspofungin, and therefore act to induce, or enhance, the previously described paradoxical growth observed at high concentration of this antifungal [35]. These data indicate that the antagonists identified do not substantially alter antifungal potency (i.e. MIC), but rather diminish the cidal capacity of caspofungin and promote tolerance. Subsequent checkerboard analysis revealed that many of the antagonists are protective at much lower concentrations than the 5 µM concentrations used in the initial screens (figure S1). Given that four of the five antagonists studied lack stand-alone antifungal activity, it was not possible to calculate FIC indices [36, 37]. However, bliss independence scores were calculated, by assessing the maximum positive Bliss deviation, which corresponds to the most antagonistic interaction with caspofungin [38] (table 2), and revealed substantial antagonistic effects over specific concentration ranges of each pairwise combination, with up to 65% growth restoration versus the minus antifungal control, by the 72-hour timepoint.

**Figure 1.**
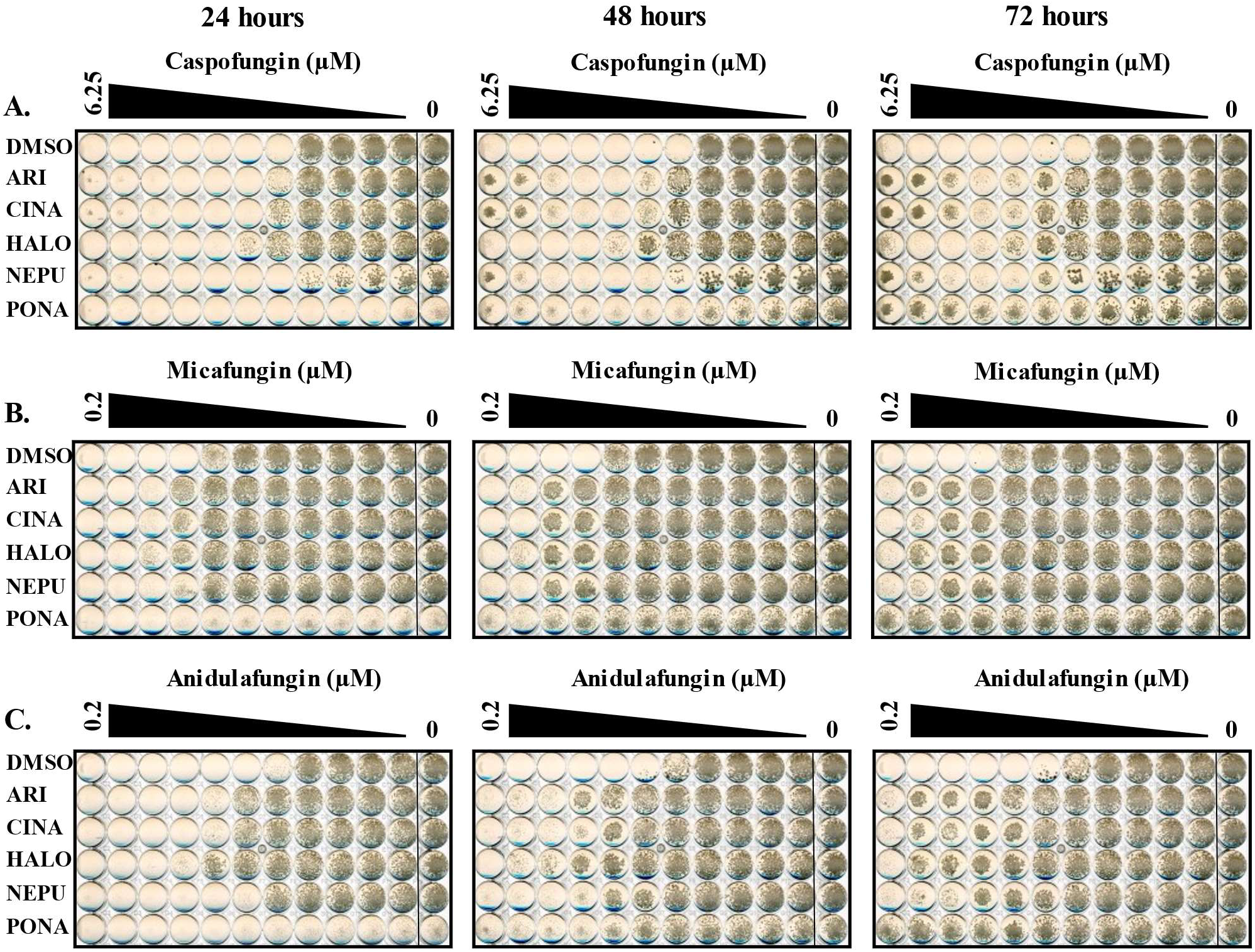
Several medications alter *Candida albicans* tolerance to multiple echinocandins. Antifungal susceptibility assays were performed with *C. albicans* SC5314. SC5314 was grown in RPMI-pH 7 (2% glucose) supplemented with two-fold dilutions of caspofungin **(A),** micafungin **(B),** or anidulafungin (C) in combination with either 5 μM of aripiprazole (ARI), 5 μM cinacalcet (CINA), 5 μM haloperidol (HALO), 5 μM netupitant (NEPU), 2.5 μM ponatinib (PONA), or vehicle (DMSO). Plates were incubated at 35°C, and imaged after 24, 48, and 72 hours. Images are representative of assays performed in biological duplicate.

**Table 2.**
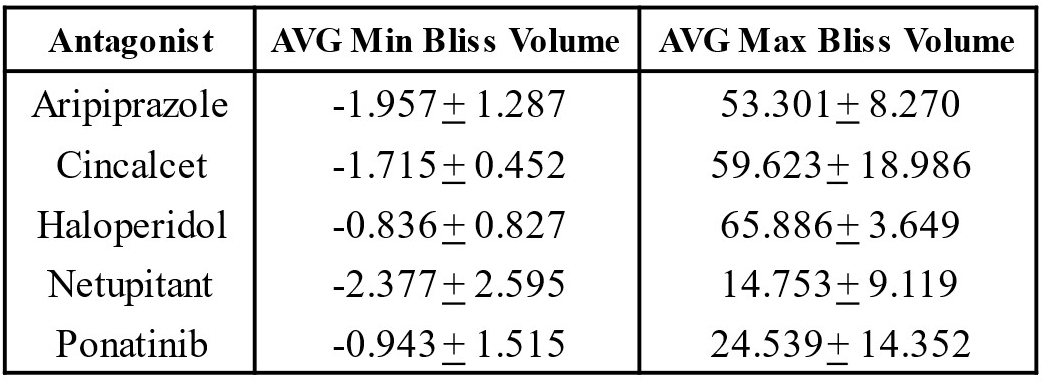
Bliss independence analysis for antagonist and caspofungin interaction. Checkerboard assays were performed using *C. albicans* SC5314 in RPMI-pH 7 (2% glucose) Plates were incubated for 72 hours (at 35°C). Bliss volume were calculated for each combination of antagonist and caspofungin. Data is representative of checkerboards performed in biological triplicate.

| Antagonist | AVG Min Bliss Volume | AVG Max Bliss Volume |
| --- | --- | --- |
| Aripiprazole | -1.957± 1.287 | 53.301± 8.270 |
| Cinacalcet | -1.715± 0.452 | 59.623± 18.986 |
| Haloperidol | -0.836± 0.827 | 65.886± 3.649 |
| Netupitant | -2.377± 2.595 | 14.753± 9.119 |
| Ponatinib | -0.943± 1.515 | 24.539± 14.352 |

All 5 of the selected drugs also enhanced SC5314 growth in the presence of supra-MIC concentrations of anidulafungin and micafungin, the two other echinocandins approved for human use (figure 1B-C). All 5 also opposed the antifungal activity of caspofungin on *C. albicans* strains TW1 [39], and ATCC10231 (figure 2A/B). However, in the case of anidulafungin, the antagonists caused an overt increase in MIC, rather than enhanced paradoxical growth. Thus, their antagonistic activity is neither strain nor echinocandin specific. Interestingly, at least 3 of the 5 antagonists (aripiprazole, cinacalcet and halofantrine) increased the concentration of caspofungin required to suppress *Candida parapsilosis* growth (figure S2A), while both cinacalcet and netupitant also enhanced caspofungin tolerance in *Candida tropicalis* (figure S2B), although the effects were less pronounced than that observed for *C. albicans*. Thus, at least a subset of the echinocandin antagonists identified affect the response of multiple medically important *Candida* species.

**Figure 2.**
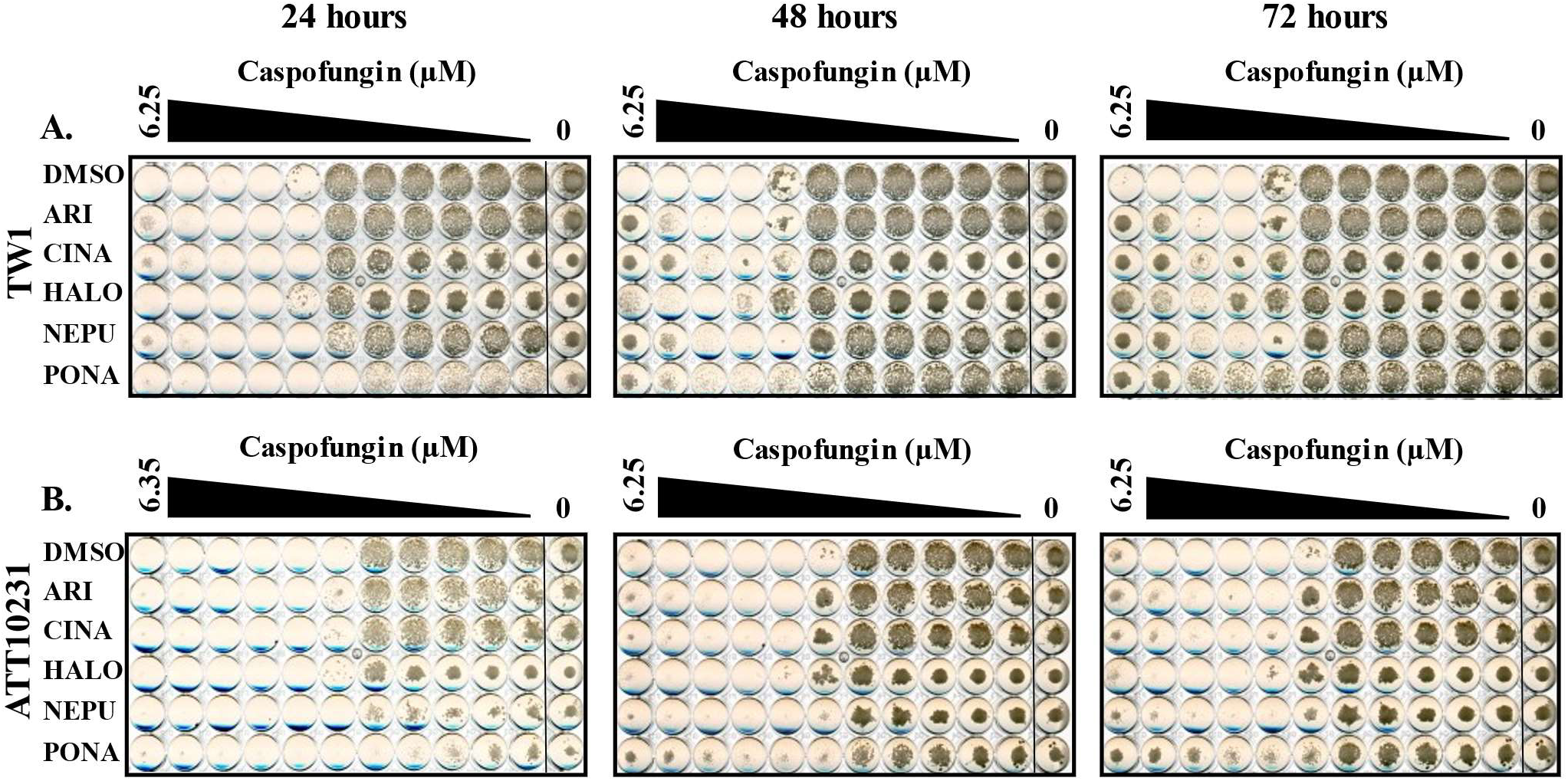
Echinocandin antagonist activity is not strain specific. *C. albicans* strain TW1 **(A)** or ATCC12031 **(B)** according to CLSI-protocol with minor revisions. Strains were grown in RPMI-pH 7 (2% glucose) supplemented with 5 μM aripiprazole (ARI), 5 μM cinacalcet (CINA), 5 μM haloperidol (HALO), 5 μM netupitant (NEPU), 2.5 μM ponatinib (PONA), or vehicle (DMSO) in combination with increasing caspofangin concentrations. Plates were incubated at 35°C, and imaged after 24, 48, and 72 hours. Images are representative of assays performed in biological duplicate.

### Antagonists diminish cidality and induce echinocandin tolerance

To further characterize the mode by which the selected antagonists protect *C. albicans* from the echinocandins activity, we conducted a more thorough assessment of cell growth, injury and viability. Growth kinetics in the presence of inhibitory concentrations of caspofungin revealed that the antagonists permit growth, but at much slower rates than in the absence of the antifungal (figure 3). Furthermore, their affect is more profound at high concentrations of the antifungal. This is consistent with them inducing a form of echinocandin tolerance rather than insensitivity, specifically, enhancing the paradoxical effect.

**Figure 3.**
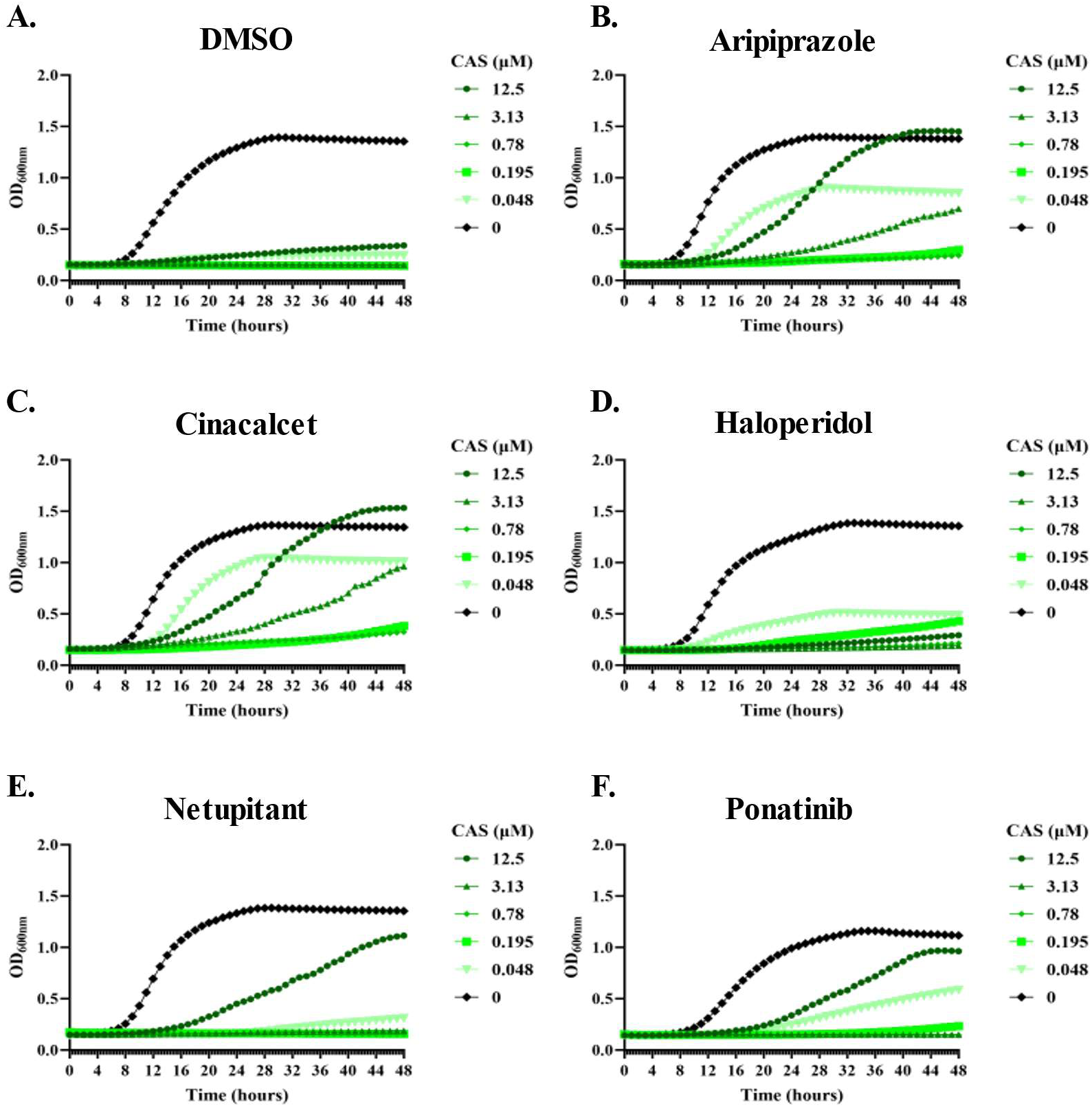
Echinocandin antagonists restore *C. albicans* growth in the presence of various caspofungin concentrations. SC5314 was grown in RPMI-pH 7 (2% glucose) medium at 35°C increasing caspofungin (CAS) concentrations in in combination with vehicle (DMSO - **A),** 5 μM aripiprazole **(B),** 5 μM cinacalcet **(C),** 5 μM haloperidol **(D),** 5 μM netupitant **(E),** or 2.5 μM ponatinib (F). Plates were incubated for 48 hours and OD_600nm_ measured every 30 minutes . Results are representative of assays performed in biological duplicate.

A *C. albicans* strain expressing a small cytoplasmic luciferase (NanoLuc™, Promega) [40] was used to compare the overall levels of damage sustained by populations of cells. Following 4-hours of caspofungin exposure at either 4 or 16X the MIC, NanoLuc release into culture supernatant was approximately 100-fold higher than mock-treated controls (figure 4A), indicating substantial levels of cell injury and/or lysis. Comparable levels of NanoLuc were released in the presence of each of the antagonistic drugs, suggesting they do not diminish the damage sustained following caspofungin exposure per se. Cell viability was also quantified as colony forming units (CFU) after 4-hours exposure to 4X MIC of caspofungin. As expected, 4X MIC of caspofungin profoundly suppressed *C. albicans* viability (figure 4B). The presence of four antagonists improved cell survival in the presence of caspofungin by approximately 10-fold, although this difference was not statistically significant due to substantial variation in CFU’s between experiments. Nonetheless, these data suggest that the antagonists protect a sub-population of *C. albicans* cells from the cidal effects of the echinocandins, and permit continued proliferation of the surviving cells at a much- reduced rate.

**Figure 4.**
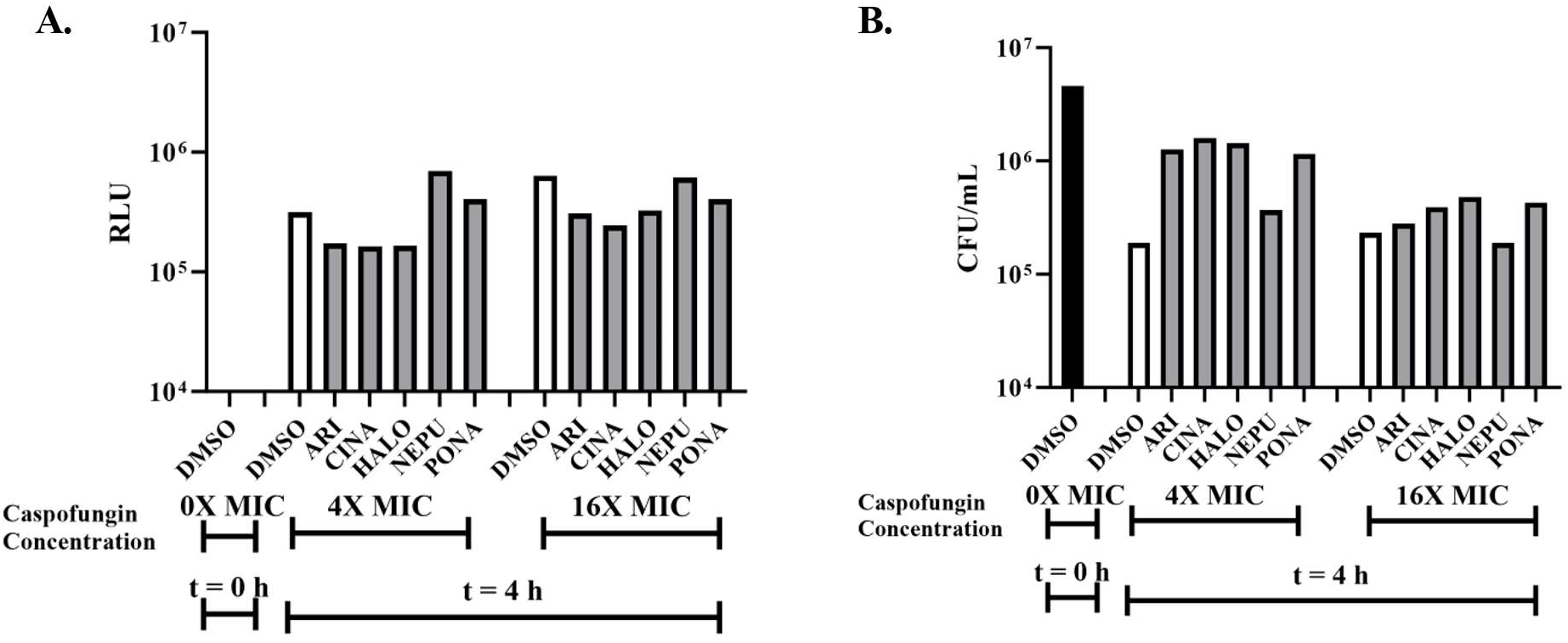
Antagonists protect *Candida albicans* viability during echinocandin exposure. **A.** *Candida albicans* wild-type GP1 and strain overexpressing NanoLuc (NLUC) were subcultured at 5 x 10^6^ cells/ml into RPMI-pH 7 (2% glucose) supplemented with vehicle (0.5% DMSO), or 5 μM aripiprazole (ARI) 5 μM cinacalcet (CINA), 5 μM haloperidol (HALO), 5 μM netupitant (NEPU), or 2.5 μM ponatinib (PONA) for 2 hours at 35°C, then caspofungin was added at 0, 4X, and 16X MIC, and further incubated for 4 hours at 35°C. Culture supernatant was collected, and NLUC release quantified for all samples, and expressed as relative luminescence units (RLU) with NLUC signal normalized to untreated controls. B. The viability of cells in each group was quantified as colony forming units (CFUs) after plating samples onto YPD agar. Representative data is shown from three independent experiments.

### Echinocandin antagonistic drugs require Mkc1p

To provide insight into the molecular mechanisms underlying echinocandin antagonism, a collection of 91 deletion mutants lacking genes encoding kinases or kinase related products were probed [29]. Initially, the collection was screened to identify gene deletion strains with caspofungin sensitivity. The first screen identified five sensitive mutants with reduced growth compared to wild-type after 48-hours in the presence of sub-inhibitory concentrations of caspofungin (0.25X MIC) (table 3 and figure 5). A further eight grew in the presence of supra-growth inhibitory concentrations (4 or 16X MIC), including mutants lacking Hog1p, as well as the Pbs2p MAPKK and Ssk2p MAPKKK that regulate Hog1p activity [41] (table 4 and figure 5). A third screen identified fifteen mutants as deficient in paradoxical growth - measured after 72-hours incubation with 64X MIC of caspofungin (table 5 and figure 5). These varied phenotypes (sensitivity as well as the capacity to survive and replicate in the presence of various concentrations of caspofungin) indicate that kinase mediated signaling is a critical determinant of *C. albicans* cell fate following echinocandin exposure, with both pro-survival and pro-cidal functions evident.

**Figure 5.**
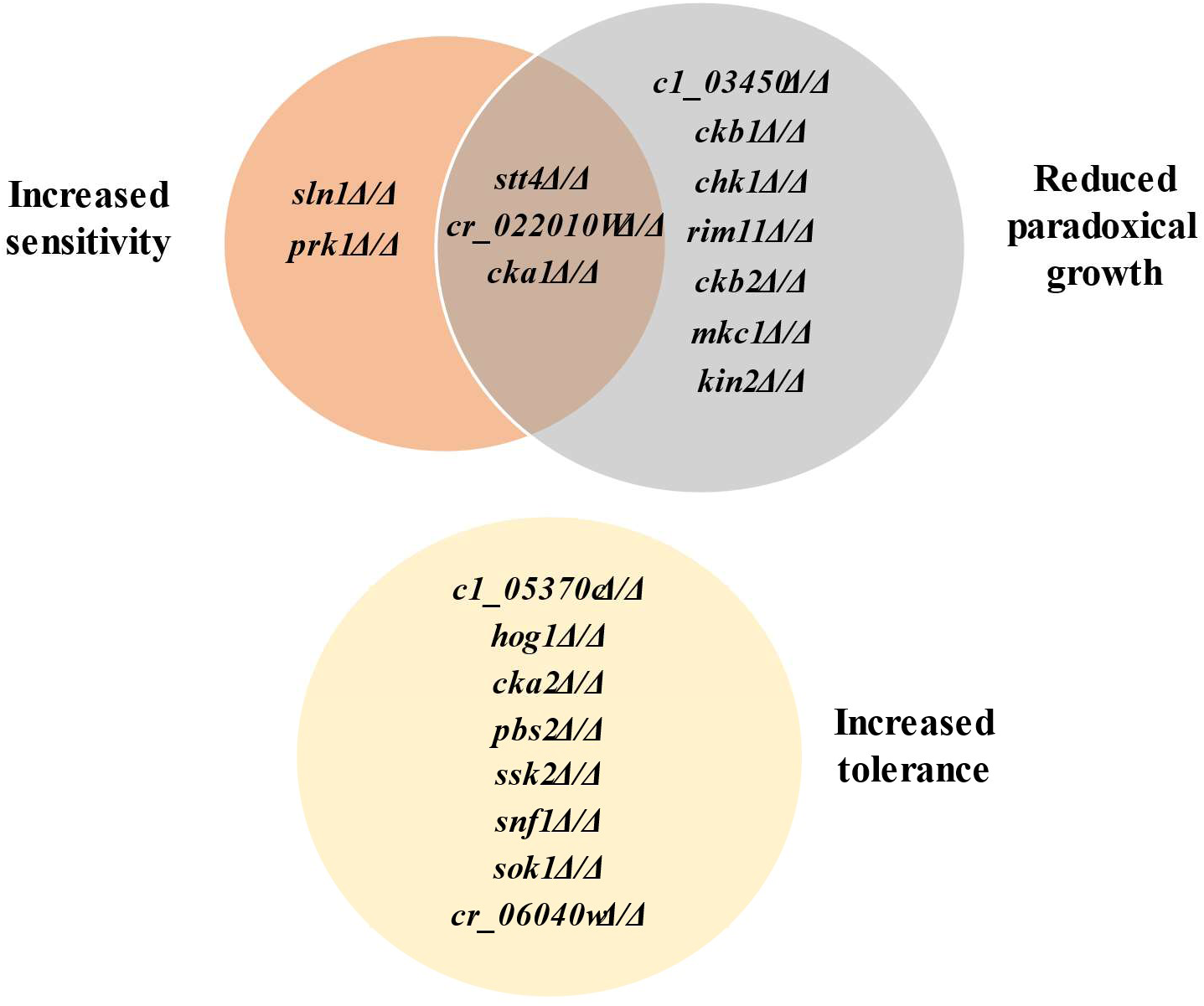
Kinase gene deletion mutants with altered echinocandin sensitivity. Ninety-one *C. albicans* deletions strains were suspended in RPMI-pH 7 (2% glucose) supplemented with 0.25X, IX, 4X, 16X, or 64X MIC caspofungin. Plates were then incubated for 72 hours at 35°C, and growth monitored at 24-hour intervals. Those that had reduced or cleared growth compared versus the wild-type control at 0.25X MIC were classified as increased sensitivity. Mutants that had a lack of paradoxical growth versus the wild-type control at 64X MIC were classified as reduced paradoxical growth. Finally, mutants with increased growth versus the wild-type control at 1,4, and 16X MIC caspofungin were identified as those with increased tolerance.

**Table 3.** List of mutants identified as caspofungin sensitive. *as described in Candida Genome Database.

| Gene name | Pathway* |
| --- | --- |
| *SLN1* | Cell wall biosynthesis |
| *PRK1* | Putative protein serine/threonine kinase |
| *cr_02210w* | Putative serine/threonine protein kinase; involved in regulation of biofilm formation |
| *STT4* | Phosphatidylinositol 4-kinase; forms a complex with Ypp1p and Efr3p that is required for phosphatidylinositol 4-phosphate, PI(4)P, in plasma membrane |
| *CKA1* | Putative alpha subunit (catalytic subunit) of protein kinase CK2 |

**Table 4.** List of mutants identified as caspofungin tolerant. *as describedin Candida Genome Database.

| Gene name | Pathway* |
| --- | --- |
| *c1_05370c* | Putative serine/threonine kinase |
| *HOG1* | MAP kinase of osmotic -, heavy metal-, and core stress response; role in regulation of response to stress |
| *CKA2* | Catalytic alpha -subunit of protein kinase CK2; interaction with calcineurin pathway |
| *PBS2* | MAPK kinase (MAPKK); role in osmotic and oxidative stress responses |
| *SSK2* | MAP kinase kinase kinase (MAPKKK); regulates Hog1 activation and signaling |
| *SNF1* | Functional homolog of *S. cerevisiae* Snf1p, which regulates sugar metabolism |
| *SOK1* | Protein kinase required for degradation of Nrg1p |
| *cr_06040w_a* | Ortholog of *S. cerevisiae* Sat4; amphotericin B induced; clade -associated gene expression |

**Table 5.** List of mutants identified as lacking paradoxical growth during caspofungin treatment. *as described in Candida Genome Database.

| Gene name | Pathway* |
| --- | --- |
| *c1_03450c* | Protein kinase-related protein, possibly involved in sterol transport between the plasma membrane and ER |
| *CKB1* | Regulatory subunit of protein kinase CK2 (casein kinase II), beta subunit |
| *CHK1* | Histidine kinase; 2-component signaling, cell wall synthesis |
| *RIM11* | Ortholog of *S. cerevisiae* Rim11p; a protein kinase involved in meiosis and sporulation in *S. cerevisiae* |
| *CKB2* | Regulatory subunit of protein kinase CK2 (casein kinase II) |
| *BCK1* | Ortholog of *S. cerevisiae* Bck1p; MAP kinase kinase kinase of cell integrity pathway |
| *MKC1* | MAP kinase; role in biofilm formation, contact-induced invasive filamentation, systemic virulence in mouse, cell wall structure/maintenance |
| *KIN2* | Protein with similarity to *S. cerevisiae* Kin2p, transcription is positively regulated by Tbf1p |
| *cr_02210w* | Putative serine/threonine protein kinase; involved in regulation of biofilm formation |
| *STT4* | Phosphatidylinositol 4-kinase; forms a complex with Ypp1p and Efr3p that is required for phosphatidylinositol 4-phosphate, PI(4)P, in plasma membrane |
| *CKA1* | Putative alpha subunit (catalytic subunit) of protein kinase CK2 |

Additional screens were performed with 4X MIC of caspofungin in the presence of each of the 5 antagonistic agents, non-responsive mutants identified (i.e. no antagonism) and follow-up dose responses conducted with caspofungin +/- antagonist to confirm phenotype. Between 8 and 13 mutants were confirmed to have no or reduced antagonistic responses for each antagonist (table 6 and figure 6). Substantial overlap was observed in the results of each screen - with a total of 10 mutants identified as non-responsive to at least 4 of the 5 antagonists tested - suggesting shared underlying mechanisms are responsible. Most conspicuous were strains lacking *BCK1*, *MKK2* and *MKC1* that encode components of the same MAP kinase signaling module, which has an established role in cell wall maintenance, caspofungin responses and induction of hyphal growth through activation of the Cph1p transcription factor [29, 42]. The importance of Cph1p was confirmed by the failure of a *cph1Δ/Δ* mutant to exhibit caspofungin antagonism with any of the 5 agents tested (figure 7). To determine if any of the antagonistic drugs affect activation of the Mkc1p pathway, extracts from SC5314 cells treated with each agent were probed with anti-phospho-Mkc1p. Aside from ponatinib, which reduced levels of phospho-Mkc1p (figure 8B/C), no major differences were apparent (data not shown). The presence of 0.25X MIC of caspofungin activated Mkc1p phosphorylation as expected (figure 8C). Ponatinib again substantially suppressed Mkc1p phosphorylation, however, the other 4 drugs had only minimal affects. Thus, while the Mkc1p pathway is necessary for echinocandin antagonism with all 5 of the selected drugs, only one of the 5 agents had clear effects on this pathway’s activity. The echinocandin antagonistic activity of 4 of the 5 drugs tested was also reduced in a *cek1Δ/Δ* mutant lacking an ERK-family protein kinase, which also has an established role in cell wall stress responses [43]. However, the affects observed were less dramatic than observed for the Mkc1p pathway deficient mutants, with the effects most obvious for ponatinib (figure 9). In addition, mutants lacking the *CST20* and *STE11* encoded kinases that function upstream of Cek1p, were unaffected (figure S3), indicating this pathway is likely less important.

**Figure 6.**
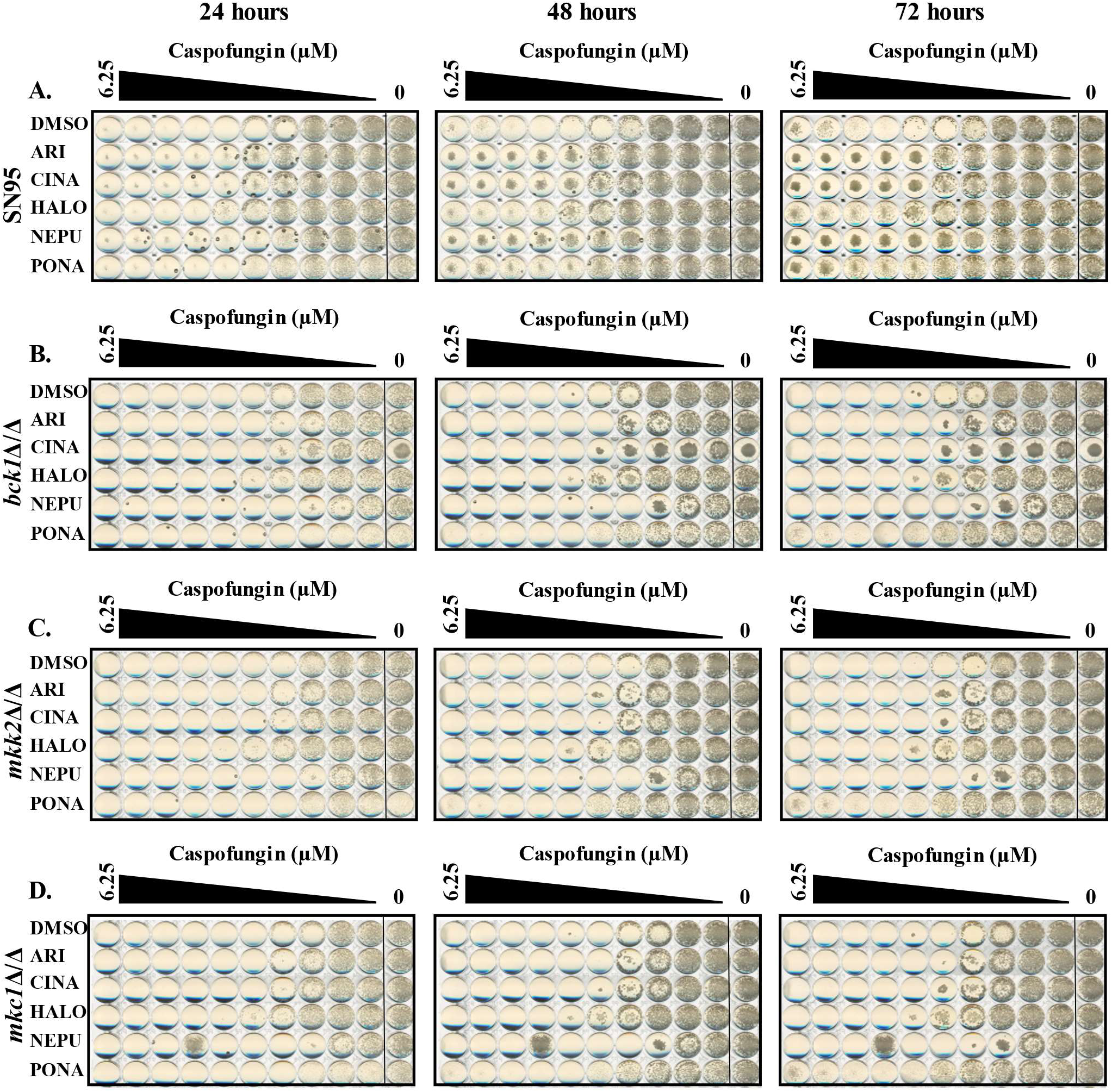
Loss of the MKC1 MAP kinase pathway function ablates echinocandin antagonism. *C. albicans* wild-type (SN95) **(A),** *hcklA/A* **(B),** *mkk2A/A* **(C),** and *mkclA/A* **(D)** mutants were grown in RPMI-pH 7 (2% glucose) supplemented with either 5 μM aripiprazole (ARI), 5 μM cinacalcet (CINA), 5 μM haloperidol (HALO), 5 μM netupitant (NEPU), 2.5 μM ponatinib (PONA), or vehicle (DMSO) and a range of caspofungin concentrations. Plates were incubated at 35°C, and imaged after 24, 48, and 72 hours. Images are representative of assays performed in biological duplicate.

**Figure 7.**
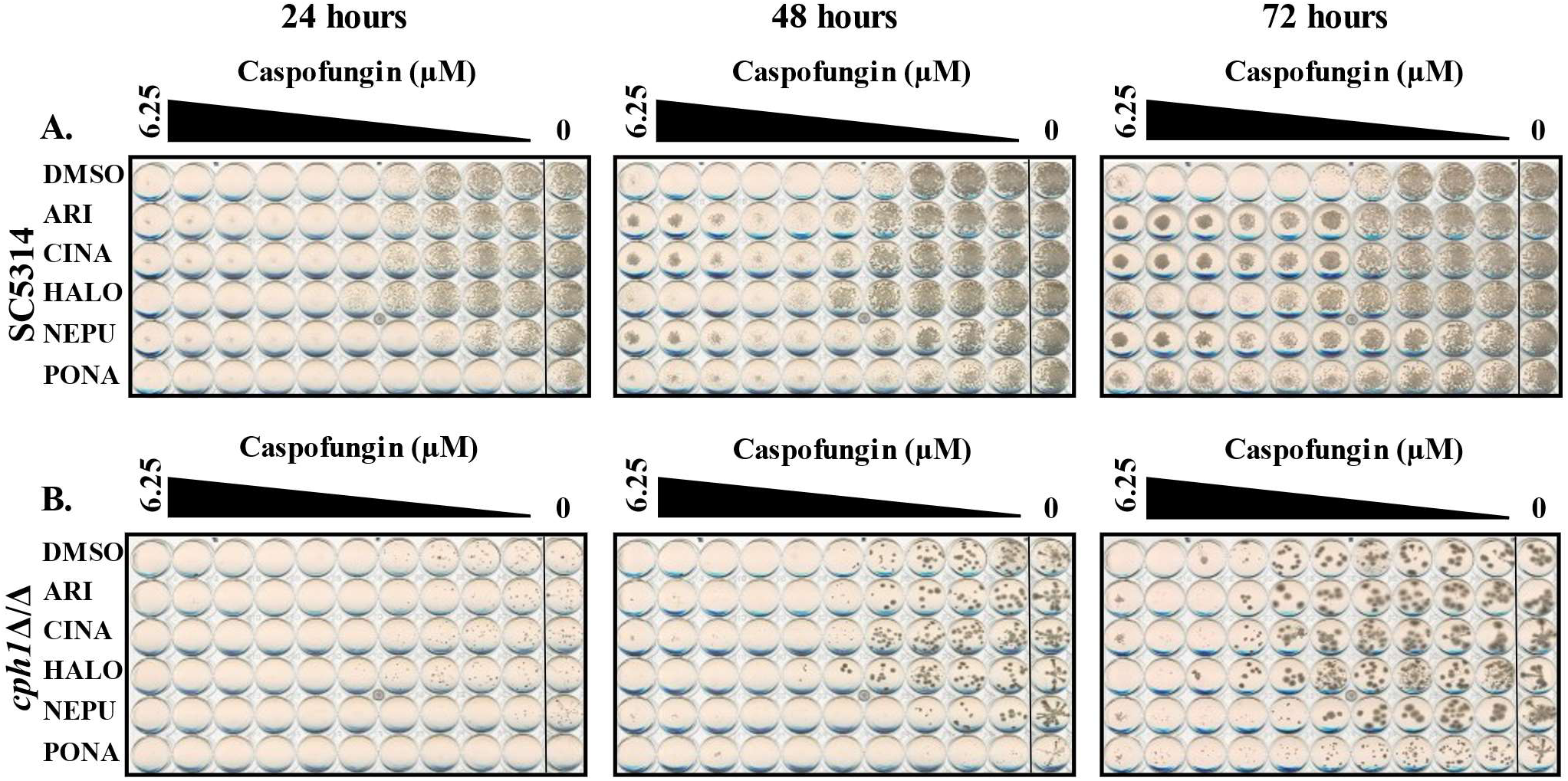
Echinocandin antagonism is lost in *C. albicans cphlA/A* mutant. *C. albicans* SC5314 **(A)** and *cphlA/A* **(B)** mutant were grown in RPMI-pH 7 (2% glucose) supplemented with either 5 μM aripiprazole (ARI), 5 μM cinacalcet (CINA), 5 μM haloperidol (HALO), 5 μM netupitant (NEPU), 2.5 μM ponatinib (PONA), or vehicle (DMSO) in combination with increasing caspofungin concentrations. Plates were incubated at 35°C, and imaged after 24, 48, and 72 hour’s. Images are representative of assays performed in biological duplicate.

**Figure 8.**
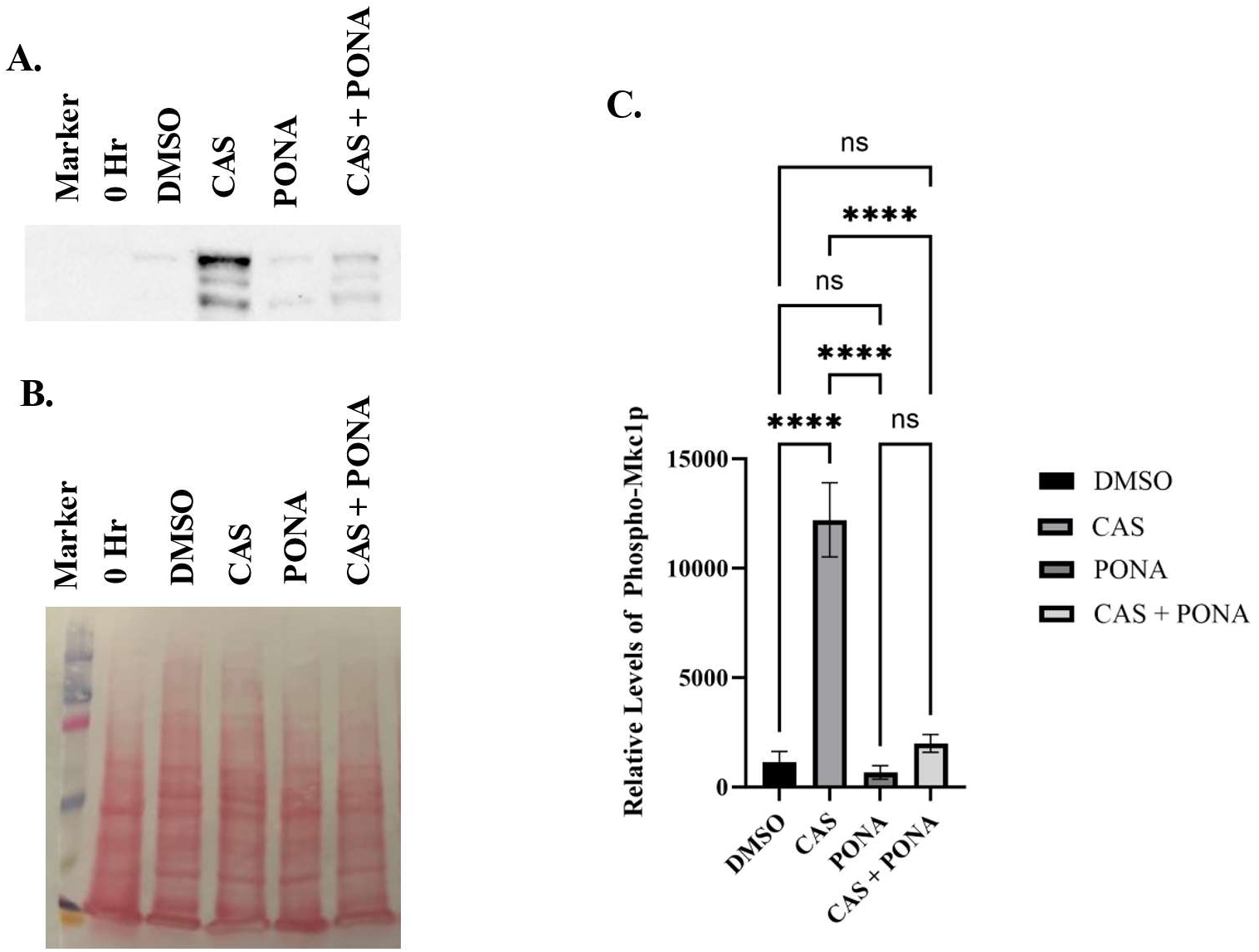
Ponatinib suppresses echinocandin induced Mkclp phosphorylation. **A-B. *C.*** *albicans* SC5314 was grown for two hours in RPMI medium supplemented with 0.5% DMSO (vehicle), 0.05 μM caspofungin (CAS), or 2.5 μM ponatinib (PONA), or both drugs in combination. Cell extracts were prepared and levels of phospho-Mklcp quantified through immunoblot analysis with anti-phospho-p44/42 antibody and anti­rabbit IgG (H+L) **(A)** compared to overall protein content **(B). C.** Quantification of the phosphorylation via densitometric analysis for four independently performed experiments. Data is the mean + standard deviation. Statistical significance was calculated via one-way ANOVA with Dunnet’s post test.

**Figure 9.**
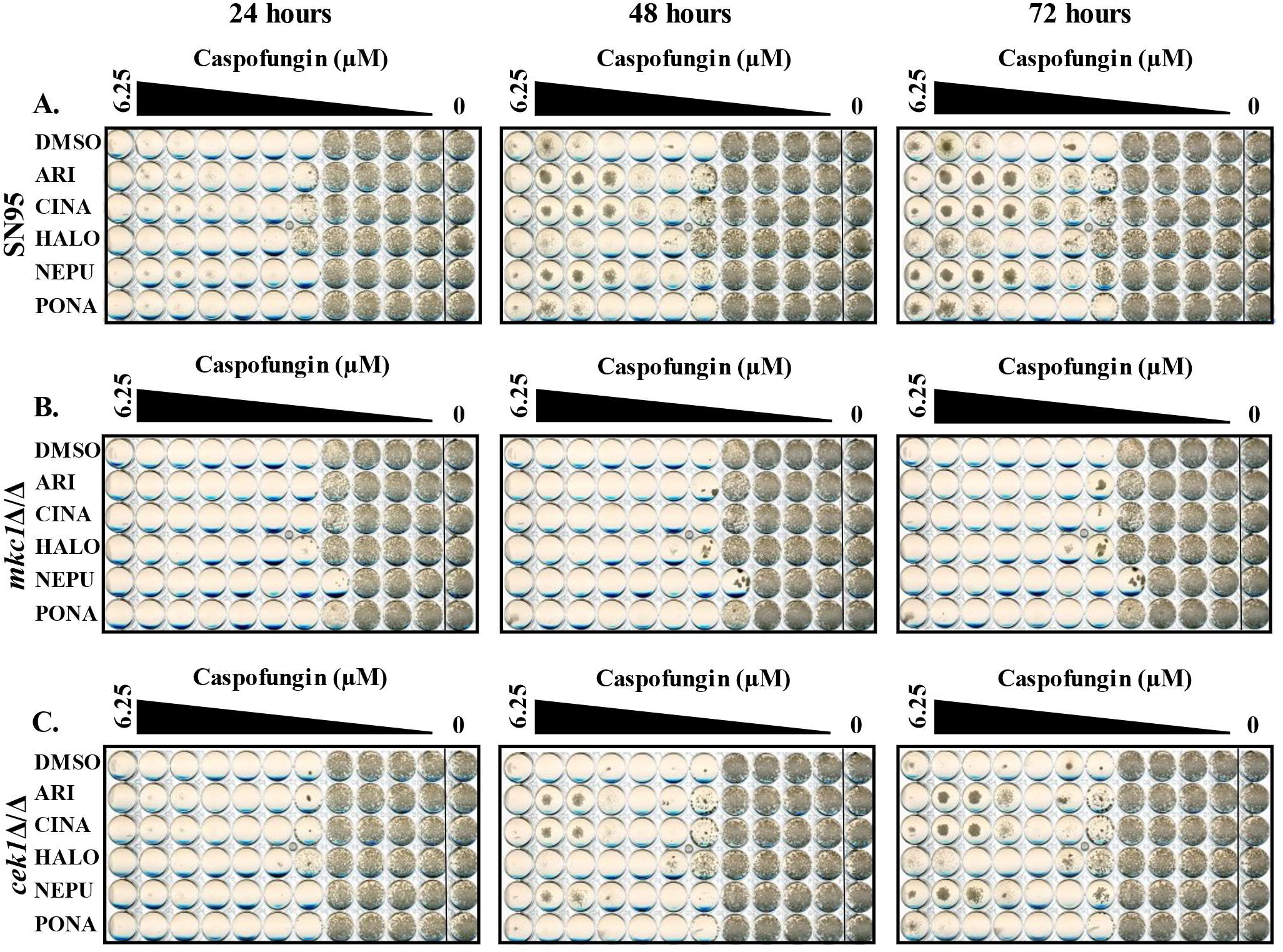
Deletion of *MKC1* eliminates the tolerance phenotype. *C. albicans* wild-type (SN95) **(A),** *mkclA/A* **(B),** and *ceklA/A* **(C)** mutants in RPMI-pH 7 (2% glucose) supplemented with either 5 μM aripiprazole (ARI), 5 μM cinacalcet (CINA), 5 μM haloperidol (HALO), 5 μM netupitant (NEPU), 2.5 μM ponatinib (PONA), or vehicle (DMSO) in combination with increasing caspofungin concentrations. Plates were incubated at 35°C, and imaged after 24, 48, and 72 hours. Images are representative of assays performed in biological duplicate.

**Table 6.** List of C. albicans mutants non-responsive to echinocandin antagonists. *as describedin Candida Genome Database.

| Gene name | Antagonists affected | Gene description* |
| --- | --- | --- |
| *SWE1* | ARI, CINA, HALO, NEPU, PONA | Putative protein kinase with a role in control of growth and morphogenesis |
| *CKB1* | ARI, CINA, HALO, NEPU, PONA | Regulatory subunit of protein kinase CK2 (casein kinase II), beta subunit |
| *CKB2* | ARI, CINA, HALO, NEPU, PONA | Regulatory subunit of protein kinase CK2 (casein kinase II), beta' subunit |
| *MKC1* | ARI, CINA, HALO, NEPU, PONA | MAMAP kinase; role in biofilm formation, contact -induced invasive filamentation, systemic virulence in mouse, cell wall structure/maintenance |
| *STT4* | ARI, CINA, HALO, NEPU, PONA | Phosphatidylinositol -4-kinase; forms a complex with Ypp1p and Efr3p that is required for phosphatidylinositol -4-phosphate, PI(4)P, in plasma membrane |
| *MKK2* | ARI, CINA, NEPU, PONA | Ortholog of *S. cerevisiae* Mkk2p; MAP kinase kinase involved in signal transduction |
| *RAD53* | ARI, CINA, HALO, PONA | Protein involved in regulation of DNA -damage-induced filamentous growth; putative component of cell cycle checkpoint |
| *CEK1* | ARI, CINA, HALO, PONA | ERK-family protein kinase; required for wild -type yeast-hypha switch, mating efficiency, virulence in mice; Cst20 -Hst7-Cek1-Cph1 MAPK pathway |
| *BCK1* | ARI, CINA, HALO, NEPU | Ortholog of *S. cerevisiae* Bck1p; MAP kinase kinase kinase of cell integrity pathway |
| *PRK1* | ARI, CINA, HALO, NEPU | Putative protein serine/threonine kinase |
| *KIN2* | ARI, CINA, HALO | Protein with similarity to *S. cerevisiae* Kin2p |
| *SAK1* | ARI, CINA | Serine/threonine protein kinase, acts as an upstream activating factor for the SNF1 complex that regulates responses to nutrient stress |
| *HS11* | ARI, CINA | Probable protein kinase involved in determination of morphology during the cell cycle of both yeast-form and hyphal cells via regulation of Swe1p and Cdc28p |

### A core set of genes associated with xenobiotic response and hyphal growth are responsive to echinocandin antagonistic drugs

To provide additional insight into the mechanisms underlying the protective activity of the five selected antagonists, as well as their overall impact upon *C. albicans* physiology, we determined how each affects global patterns of gene transcription. SC5314 was grown at 35°C in RPMI medium supplemented with 5 µM of each antagonist (2.5 µM for ponatinib, which is growth inhibitory at 5 µM), or 0.5% DMSO (vehicle control), total RNA extracted and analyzed by high-throughput sequencing. Antagonist-responsive genes were defined as those in which the relative transcript read frequency was increased or decreased by ≥ 2-fold (P < 0.05) in the presence of drug versus vehicle control, in each of 2 independent experiments. The number of responsive genes for each individual antagonist ranged between 590 and 42, with several transcripts affected by multiple drugs - indicating at least some similarities in their affects upon *C. albicans* physiology. We identified 17 transcripts as increased in response to at least four of the five antagonists tested, 7 of which are associated with fluconazole-resistance or with regulation by the Tac1p transcription factor (table 7, table S1), including *CDR2* which encodes an ATP-driven multi-drug efflux pump. An additional 10 of 43 transcripts elevated by any three of the five drugs, were also associated with response or sensitivity to the azole antifungals, or Tac1p dependent regulation, including that of *CDR1*, encoding a second ATP-driven multi-drug efflux pump. Of the 22 gene transcripts repressed in the presence of at least four of the antagonists, 9 are either required for *C. albicans* hyphal growth or exhibit hyphal specific expression, including that of *ECE1* – which encodes candidalysin [44]; *ALS3* and *HWP1* – both hyphal cell wall associated adhesins that mediate attachment to mammalian cells [45, 46]; and *UME6* – a transcription factor that promotes sustained hyphal growth [47]. A further 11 of 41 transcripts downregulated by any three antagonists were also associated in some way with hyphal growth, including *CHT2* encoding a chitinase previously shown to be suppressed following echinocandin exposure [48]. Many of these morphogenesis related genes are known to be induced following activation of the Ras-cAMP-PkA-Efg1p based signaling module, which plays a major role in the regulation of the yeast-hypha transition [49–51]. Additionally, two transcripts previously identified as the most responsive to loss of cAMP production [52] were also affected, with *ASR1* induced by aripiprazole, cinacalcet and netupitant, and suppressed by ponatinib; and *ASR2* induced by aripiprazole, cinacalcet, netupitant and ponatinib. Curiously, the elevated transcription of the *ASR1* and *ASR2* genes is indicative of enhanced cAMP signaling, while suppression of the *ECE1*, *HWP1*, *ALS3* and several other gene transcripts is consistent with diminished signaling through this pathway.

**Table 7.** List of *C. albicans* gene transcripts responsive to echinocandin antagonists. *as described in Candida Genome Database. *non applicable (NA) where there was no significant change in transcript regulation compared to vehicle treated cells alone.

| Gene name | Log<sub>2</sub> of Average Fold Change |  |  |  |  |
| --- | --- | --- | --- | --- | --- |
|  | Aripiprazole | Cinacalcet | Haloperidol | Netupitant | Ponatinib |
| *RTA3* | 3.423 | 3.706 | 1.818 | 4.198 | 2.059 |
| *WH11* | 2.809 | 3.507 | 1.984 | 3.878 | 2.661 |
| *PGA26* | -2.107 | -2.912 | -3.453 | -2.580 | -2.667 |
| *ALS3* | -2.545 | -4.114 | -4.510 | -4.839 | -1.146 |
| *SOU2* | 3.336 | 3.436 | 3.420 | 2.476 | NA |
| *orf19.6502* | 2.249 | 2.483 | 1.961 | 2.382 | NA |
| *RSN1* | 1.684 | 1.663 | 2.670 | 1.465 | NA |
| *GCY1* | 1.436 | 1.419 | 1.719 | 1.389 | NA |
| *LTV1* | -1.660 | -1.542 | -1.349 | -1.237 | NA |
| *orf19.5710* | -1.232 | -1.617 | -1.369 | -1.737 | NA |
| *SEO1* | -1.889 | -2.155 | -1.872 | -2.312 | NA |
| *orf19.3475* | -1.724 | -1.847 | -3.245 | -1.588 | NA |
| *DEF1* | -2.213 | -2.071 | -1.843 | -2.361 | NA |
| *UME6* | -2.695 | -3.967 | -2.154 | -4.213 | NA |
| *IHD1* | -3.511 | -5.346 | -2.929 | -4.296 | NA |
| *ECE1* | -3.211 | -5.788 | -2.523 | -5.915 | NA |
| *HWP1* | -3.118 | -5.969 | -2.933 | -5.876 | NA |
| *CFL11* | -4.302 | -4.847 | -4.012 | -4.788 | NA |
| *CDR2* | 5.776 | 6.432 | NA | 6.186 | 4.105 |
| *orf19.4886* | 2.527 | 3.187 | NA | 2.959 | 2.313 |
| *OYE23* | 1.828 | 2.208 | NA | 2.751 | 2.596 |
| *ASR2* | 2.287 | 2.772 | NA | 3.218 | 1.074 |
| *orf19.4216* | 2.077 | 2.556 | NA | 2.254 | 1.857 |
| *orf19.86* | 2.034 | 2.093 | NA | 2.503 | 1.654 |
| *HSP12* | 1.881 | 2.355 | NA | 2.131 | 1.565 |
| *orf19.4907* | 1.927 | 1.987 | NA | 2.438 | 1.227 |
| *IFE1* | 1.498 | 1.855 | NA | 2.240 | 1.880 |
| *SOD5* | -1.790 | -2.456 | NA | -2.170 | -2.946 |
| *orf19.6200* | -3.348 | -3.614 | NA | -4.799 | -3.497 |
| *orf19.3406* | NA | -1.193 | -1.382 | -1.206 | -1.811 |
| *PHO89* | NA | -2.685 | -2.514 | -2.941 | -1.850 |
| *GIT1* | NA | -2.433 | -3.220 | -2.046 | -2.393 |

### Inactivation of the cAMP-PkA responsive transcription factor Efg1p leads to echinocandin tolerance

Given the impact of the echinocandin antagonists upon morphogenesis responsive transcripts, the influence of each upon *C. albicans* hyphal growth was examined. All five echinocandin antagonists suppressed *C. albicans* hyphal growth on either 10% FBS or M199 agar to varying extents (figure 10A/B), with haloperidol and ponatinib having the greatest affect. In liquid RPMI medium at 35°C, aripiprazole moderately suppressed hyphal formation, as we have previously reported [34], reducing the proportion of cells forming hyphae as well as the length of those that form. Haloperidol and ponatinib also significantly reduced hyphal length under these conditions (figure 10C), while netupitant and cinacalcet had no obvious effect on cell morphology under these conditions. To further probe the connection between the cAMP-PkA-Efg1p morphogenesis related signaling pathway and echinocandin antagonism, an *efg1Δ/Δ* mutant was utilized. Dose-response experiments revealed that the *efg1Δ/Δ* mutant had an MIC of caspofungin approximately 8-fold higher, as well as elevated levels of paradoxical growth compared to wild-type that was unaffected by any of the aforementioned 5 antagonists (figure 11). The mutant also exhibited substantial levels of tolerance to both anidulafungin and micafungin (data not shown). These data indicate that the morphogenesis related transcription factor Efg1p confers susceptibility to caspofungin.

**Figure 10.**
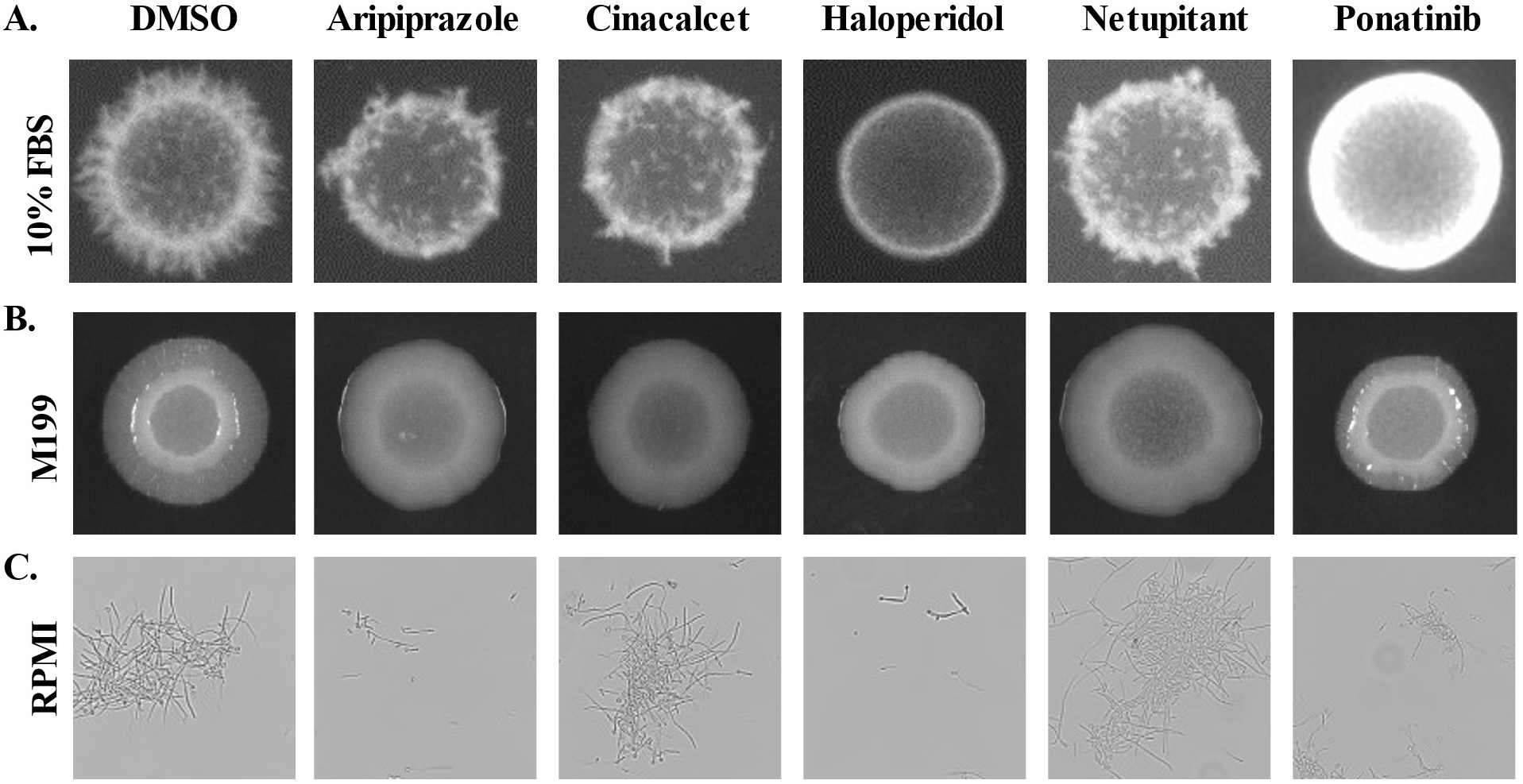
Several e chinocandin antagonists suppress *C. albicans* hyphal growth. **A-B.** SC5314 was suspended at 2 x 10^6^ cells ml and 2 pl spotted to 10% fetal bovine serum (FBS - **A)** or Ml99 (B) agar supplemented with either 5 μM aripiprazole, 5 μM haloperidol, 5 μM netupitant, 5 μM cinacalcet, 2.5 μM ponatinib, or vehicle (DMSO). Colonies were then imaged after 96 hours incubating at 37°C. C. SC5314 was suspended in liquid RPMI medium supplemented with same drug concentrations as in **(A-B).** After 6 hours of incubation, cells were fixed in 10% formalin, cell morphology was observed microscopically. Images are representative of assays performed in biological triplicate.

**Figure 11.**
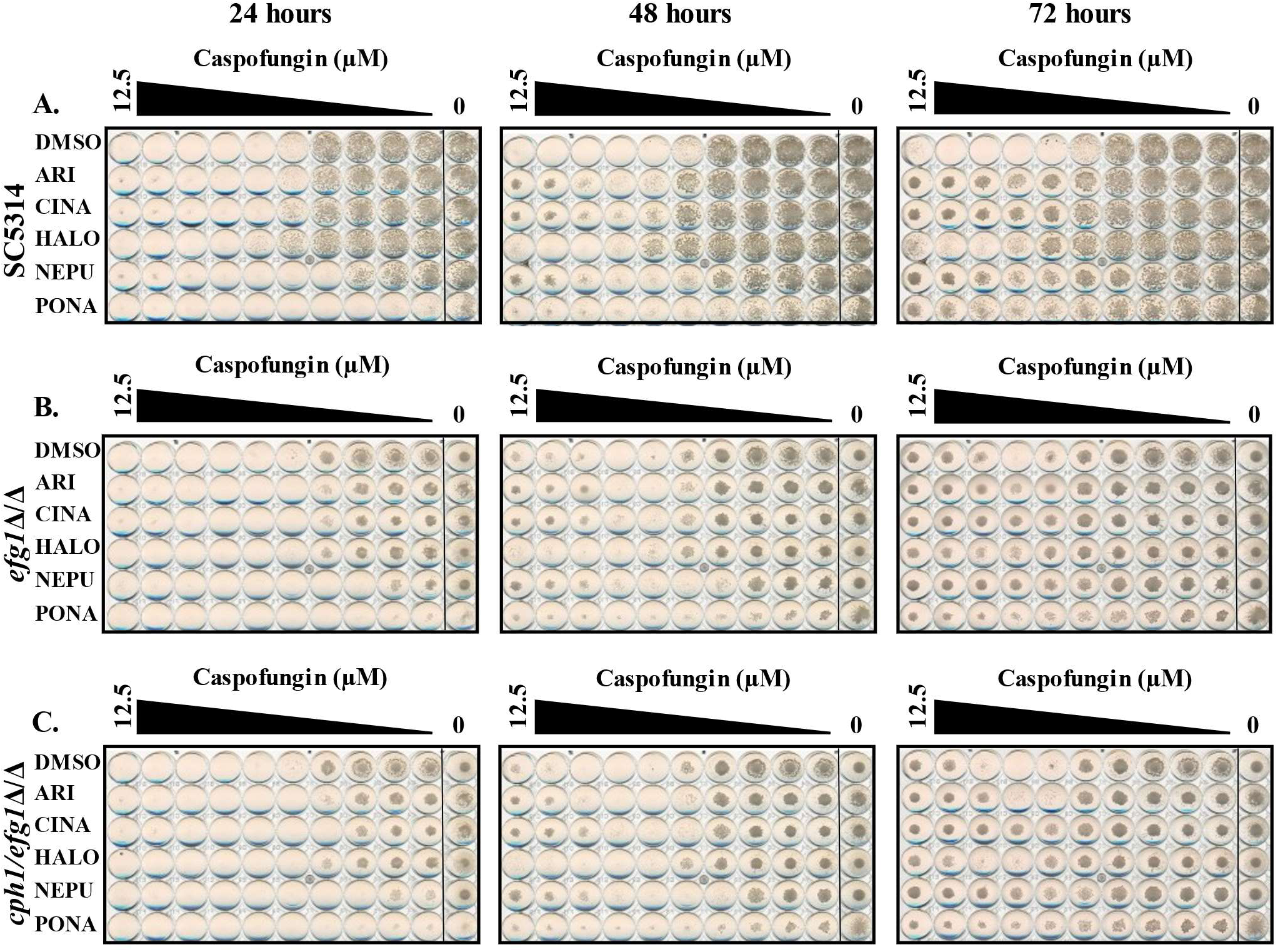
Deletion of *EFG1* or double deletion of *EFGUCPH1* does not impact echinocandin tolerance phenotype. *C. albicans* SC5314 **(A)** *efglA/A* **(B)** and *efglA/A/cphlA/A* mutant **(C)** were grown in RPMI-pH 7 (2% glucose) supplemented with either 5 μM aripiprazole (ARI), 5 μM cinacalcet (CINA), 5 μM haloperidol (HALO), 5 μM netupitant (NEPU), 2.5 μM ponatinib (PONA), or vehicle (DMSO) in combination with increasing caspofungin concentrations. Plates were incubated at 35°C, and imaged after 24, 48, and 72 hours. Images are representative of assays performed in biological duplicate.

## Discussion

The studies described herein have revealed a substantial number of drug-fungus interactions that directly or indirectly affect *C. albicans* susceptibility to the ordinarily cidal echinocandin antifungals. Several points are notable: 1. The MIC determined at the 24-hour timepoint was not (in most cases) significantly altered by the antagonistic drug, but rather residual growth observed above the MIC at 48- and 72-hours was dramatically enhanced. Based on this phenotype, as well as viability counts following caspofungin exposure, we have classified the phenomenon as a form of tolerance that promotes survival of the fungal cells, rather than resistance. For 4 of the 5 antagonists examined in detail, this also appeared consistent with an enhanced form of the paradoxical effect previously described at high concentrations of the echinocandins. However, paradoxical growth is principally observed with caspofungin, less commonly with micafungin or anidulafungin [35, 53], and is more apparent in some *C. albicans* isolates over others. In contrast, the antagonistic activities we have described affect *C. albicans* response to all three echinocandins, and their activity does not seem restricted to specific isolates. Thus, it remains uncertain how the drug-induced echinocandin antagonism observed within this study is qualitatively related to the previously described paradoxical phenotype.

Our findings suggest that the antagonism observed depends upon a complex response requiring multiple signaling pathways that have been fairly well characterized with respect to their roles in regulating yeast- hypha morphogenesis, and *C. albicans* response to cell wall stress [43, 54, 55]. Specifically, our data indicate that the Mkc1p-MAPK signaling module, which acts upon the Cph1p transcription factor [56], is crucial for the echinocandin tolerance induced by the antagonists. *CKB1* and *CKB2*, both encoding regulatory subunits of casein kinase 2, were also identified as non-responsive to the caspofungin antagonists. Curiously, we had also identified the two mutants lacking either of the two catalytic A subunits of CK2 as having opposing phenotypes, with the *cka1Δ/Δ* mutant hypersensitive to caspofungin and the *cka2Δ/Δ* mutant more tolerant than wild-type (data not shown). Signaling through the calcium- dependent calcineurin (CN) phosphatase pathway is known to be important for fungal survival upon exposure to both azole and echinocandin antifungals [57]. Blocking calcineurin (CN) signaling with either cyclosporin A or FK596 completely eliminated the antagonistic activity of all five drugs (figure S4), and the echinocandin sensitivity of a *C. albicans cnb1Δ/Δ* mutant, lacking the B regulatory subunit of CN, was unaffected by any of the 5 antagonists tested (figure S5). In contrast, 4 of the 5 antagonists (except haloperidol) promoted caspofungin tolerance in a *crz1Δ/Δ* mutant (figure S5), lacking one of the major CN-responsive transcription factors [58]. Thus, Crz1p-independent CN-signaling is also required for echinocandin antagonism. Curiously, we observed that a mutant lacking the Hog1p MAP kinase exhibited significant levels of residual growth at all concentrations of caspofungin above the MIC, that are apparently further enhanced by aripiprazole, cinacalcet, netupitant or ponatinib (figure S6). Together, these results indicate that the echinocandin antagonists tested act through a complex mechanism that requires both Mkc1p, CK2 and Crz1p-independent functions of CN. The similarities in the pathway requirements of the five antagonists studied in detail indicates a largely shared molecular mechanism with respect to antifungal antagonism, which is somewhat surprising given the diversity in their chemical structure and targets within the mammalian host. The capacity to sense these unrelated small molecules and produce a seemingly common functional response with respect to the tolerance phenotype is curious and suggests the presence of xenobiotic sensing modules and/or receptors capable of promiscuous interactions with small molecules. Indeed, analysis of transcriptional responses indicated that a significant number of echinocandin antagonists affect the transcription Tac1p-responsive genes. The transcription factor Tac1p, and close relatives from other species, have been shown to directly engage and be activated by several medically important molecules, most notably fluphenazine [59]. In a related study, we also recently described a that substantial proportion of drugs identified as fluconazole antagonists, required Tac1p for activity [28], suggesting this response module is able to sense a much broader array of molecular species than previously appreciated. However, in contrast to the fluconazole antagonists, most of which required Tac1p for activity, Tac1p is not required for the echinocandin antagonism described herein (figure S7), and its activation is therefore likely incidental. As such, Tac1p activation seems to define a more general response to encountering xenobiotics. Nonetheless, this provides clear precedence for the existence of systems within *C. albicans* that function to detect and respond to an impressive variety of molecular entities. Defining the precise molecular targets or receptors of the echinocandin antagonists that activate the protective responses is likely to be a non-trivial undertaking.

The suppression of hyphal associated gene transcription by the echinocandin antagonists was unexpected and raises questions of how cellular morphotype may affect *C. albicans* echinocandin sensitivity. The culture conditions used for the antagonism screens, as well as for the RNAseq analysis (RPMI pH 7) stimulates *C. albicans* hyphal growth. Future work should explore if modulation of yeast-hypha morphogenesis can provide a strategy for *C. albicans* to mitigate the consequences of echinocandin exposure and determine if yeast and hyphal forms are similarly susceptible to echinocandins induced cell death.

A major rationale for conducting the antagonism screen, was to identify factors that can help explain the often-unpredictable response of individual patients to echinocandin therapy i.e. could patients treated with one or more antagonistic medication be less responsive to antifungal treatment, and/or more prone to recurrence? In a broader context, this study raises questions of how biologically active molecules including the medications consumed by colonized or infected individuals affect fungal physiology, and thus potentially the equilibrium of the host-pathogen interaction. Drug-fungus interactions for example could alter the outcome of infection by: 1). Causing profound physiological dysfunction that compromises fungal viability, fitness, or pathogenicity; 2). Changing the surface characteristics of the fungus in ways that alter its immunogenicity, binding/activation of complement, or sensitivity to host derived antimicrobial peptides; or 3). Altering fungal sensitivity to antifungal drugs, as is the focus of this study. However, the highly variable, often transient and combinatorial nature of medications usually provided to patients at risk of IFIs, specific clinical circumstances (including the nature and severity of immune dysfunction), site of infection, causative species and potentially genetic factors of both the host and pathogen are likely to make it difficult, if not impossible to discern the impact of any individual drug upon antifungal efficacy based on clinical observations alone. These factors are also likely to provide significant constraints on the power of clinical studies to reliably detect the impact of individual medications upon therapeutic efficacy, or the outcome of infection, especially given the relatively small numbers of infected patients that may be treated with a given antagonistic medication. Nonetheless, these limitations do not mean that such drug-fungus interactions have no impact, and it is possible they may help explain the often- poor correlation between the results of *in vitro* antifungal susceptibility tests and clinical efficacy. A more reliable strategy may be to use animal models of infection that enable control of many of the confounding factors.

Several key factors are likely to determine if the antagonistic agents identified within this this study affect echinocandin efficacy within mammalian tissue. First, it is important to consider the effects of each drug at therapeutically relevant drug concentrations - which in the context of disseminated infection - most investigators will take to be unbound blood serum concentrations from patients on a standard dosing regimen. However, it is the available concentration of the antifungal as well as the antagonistic agent at the site of infection that is most relevant, and tissue concentrations can differ markedly from unbound serum concentrations. Additionally, acute inflammation, tissue dissolution and protein digestion by fungal enzymes within foci of infection could have further affects, and drug concentrations at site of fungal infection are unknown. Finally, *Candida* species naturally colonize the GI tract, where the concentration of orally administered drugs (or those excreted via the GI tract) is potentially much higher.

## Acknowledgements

Research reported in this publication was supported by the National Institute of Allergy and Infectious Diseases of the National Institutes of Health under Award Number R01AI152067 (to GEP). The content is solely the responsibility of the authors and does not necessarily represent the official views of the National Institutes of Health. We thank Promega Corporation for permission to produce and to utilize the *C. albicans* adapted NanoLuc coding sequence.

---

**Figure SI.**
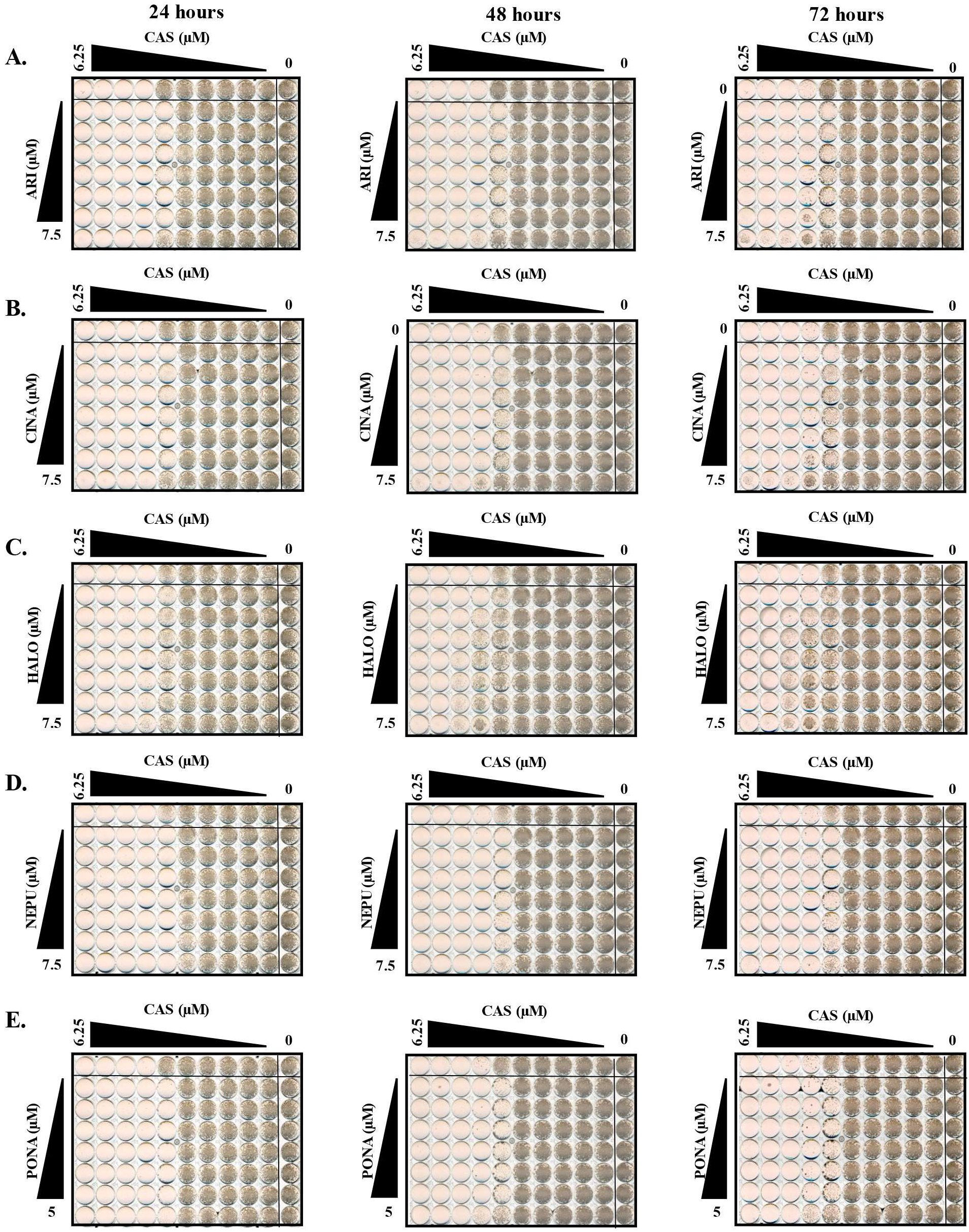
Echinocandin antagonists possess activity at sub micromotor activity. **Checkerboards assays** were performed using C. *albicans.* SC5314 was grown in RPMI-pH 7 (2% glucose) with increasing caspofungin (CAS) concentration with increasing concentrations of aripiprazole (ARI - **A),** cinacalcet (CINA - **B)** haloperidol (HALO - **C),** netupitant (NEPU - **D),** or ponatinib (PONA - **E).** Plates were incubated for up to 72 hours of incubation at 35°C, with plates imaged every 24 hours to monitor changes in tolerance. Images are representative of assays performed in biological triplicate.

**Figure S2.**
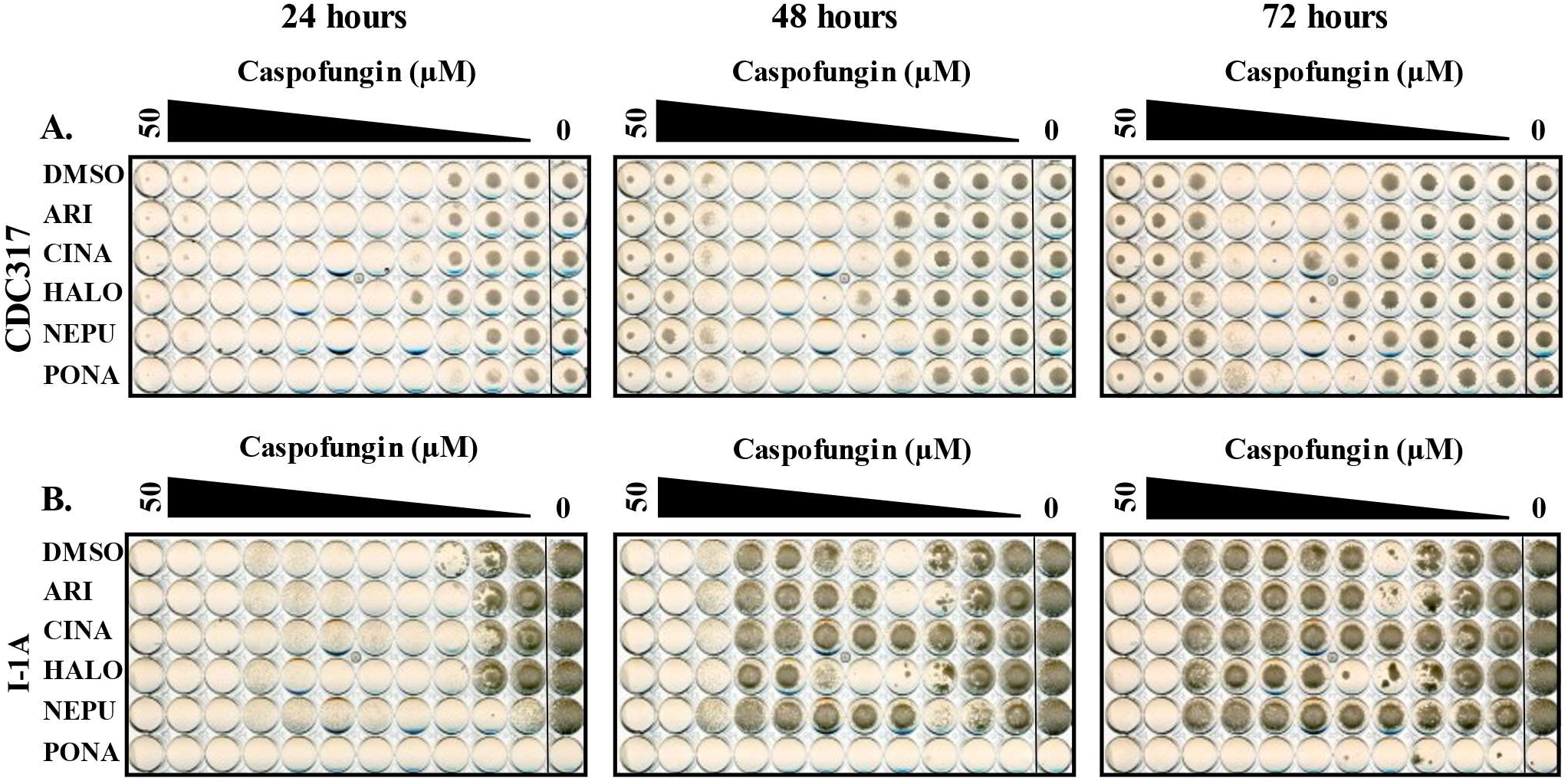
Selected antagonist activity is not species specific. *C. parapsilosis* strain **CDC317 (A)** or *C. tropicalis* strain I-1A (B) were grown in RPMI-pH 7 (2% glucose) supplemented with either 5 μM aripiprazole (ARI), 5 μM cinacalcet (CINA), 5 μM haloperidol (HALO), 5 μM netupitant (NEPU), 2.5 μM ponatinib (PONA), or vehicle (DMSO) in combination with increasing caspofungin concentrations. Plates were incubated at 35°C, and imaged after 24, 48, and 72 hours. Images are representative of assays performed in biological duplicate.

**Figure S3.**
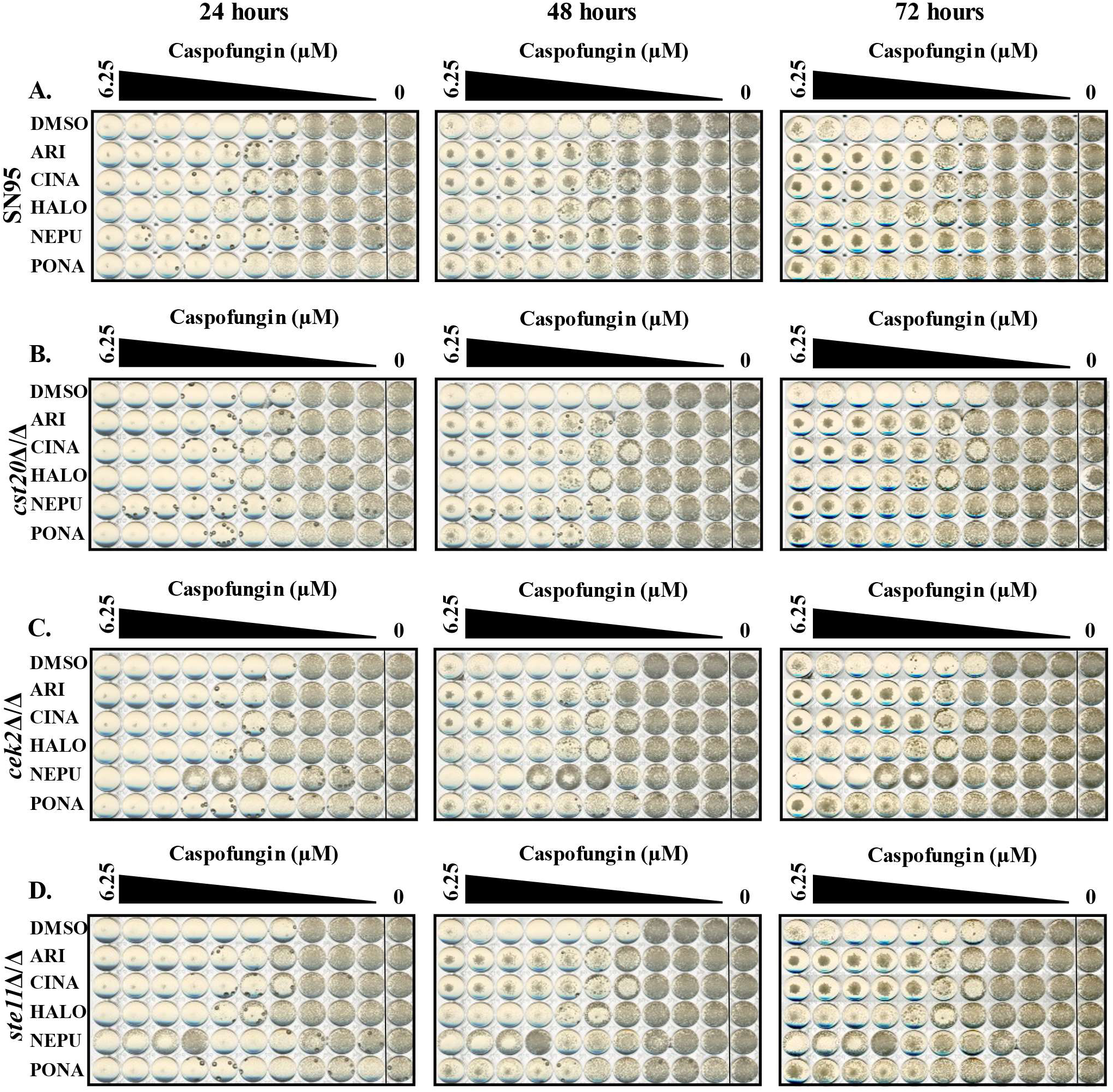
Echiuocandin antagonist activity is not dependent other components of Ceklp pathway. *C. albicans* SC5314 **(A),** *cst20A/A* **(B),** *cek2A/A* **(C),** and *stellA/A* **(D)** mutants were grown in RPMI-pH 7 (2% glucose) supplemented with either 5 μM aripiprazole (ARI), 5 μM cinacalcet (CINA), 5 μM haloperidol (HALO), 5 μM netupitant (NEPU), 2.5 μM ponatinib (PONA), or vehicle (DMSO) in combination with increasing caspofungin concentrations. Plates were incubated at 3 5 °C, and imaged after 24, 48, and 72 hours. Images are representative of assays performed in biological duplicate.

**Figure S4.**
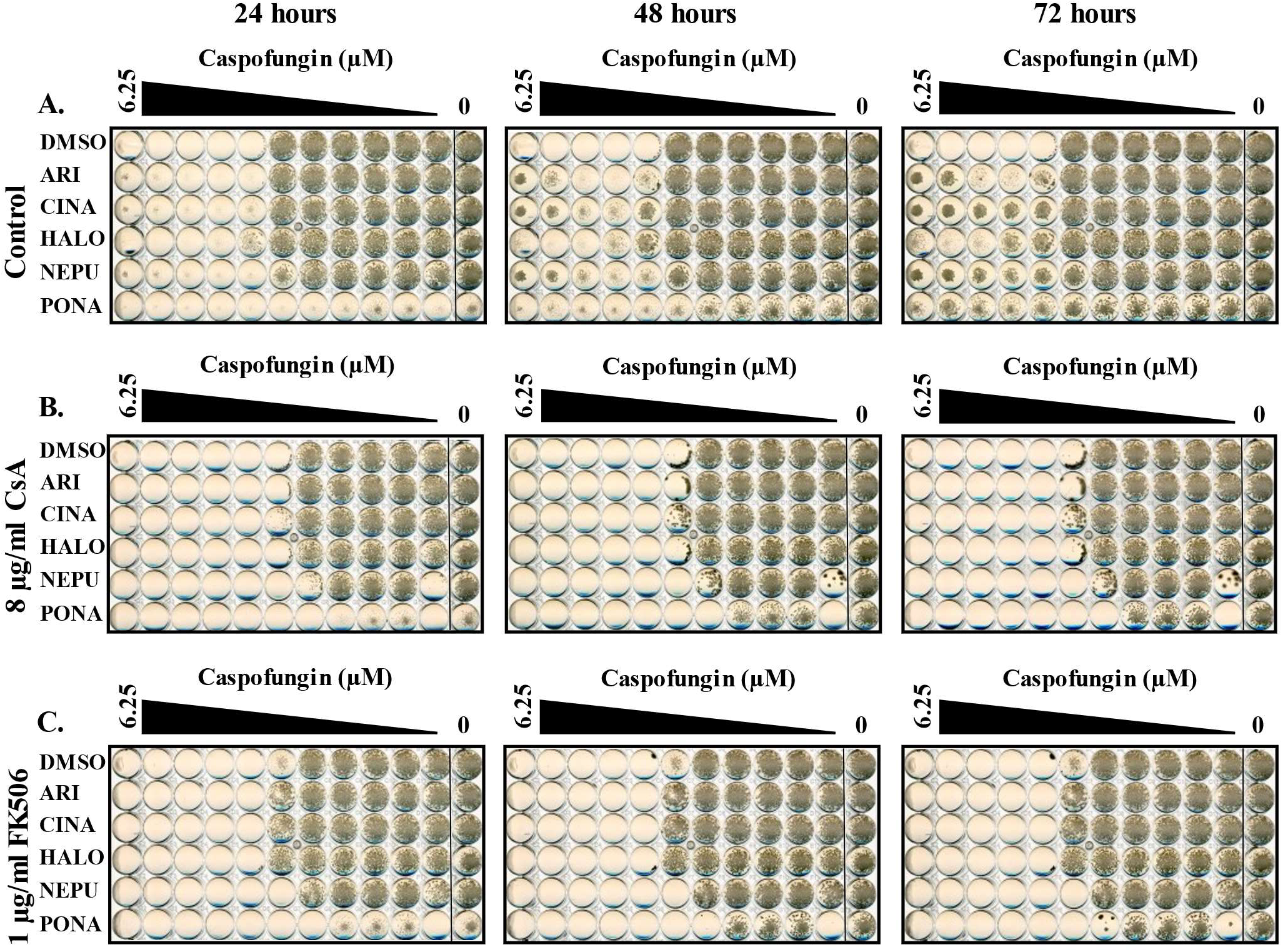
Inhibition of the calcineurin pathway suppress echinocandin antagonism. *C. albicans* SC5314 was grown in RPMI-pH 7 (2% glucose) supplemented with either 5 μM aripiprazole (ARI), 5 μM cinacalcet (CINA), 5 μM haloperidol (HALO), 5 μM netupitant (NEPU), 2.5 μM ponatinib (PONA), or vehicle (DMSO) in combination with increasing caspofungin concentrations. Medium was supplemented either with H_2_O (control - **A),** 8 pg/ml Cyclosporin A (CsA - **B),** or 1 pg/ml FK506 (C) Plates were incubated at 35°C, and imaged after 24, 48, and 72 hours. Images are representative of assays performed in biological duplicate.

**Figure S5.**
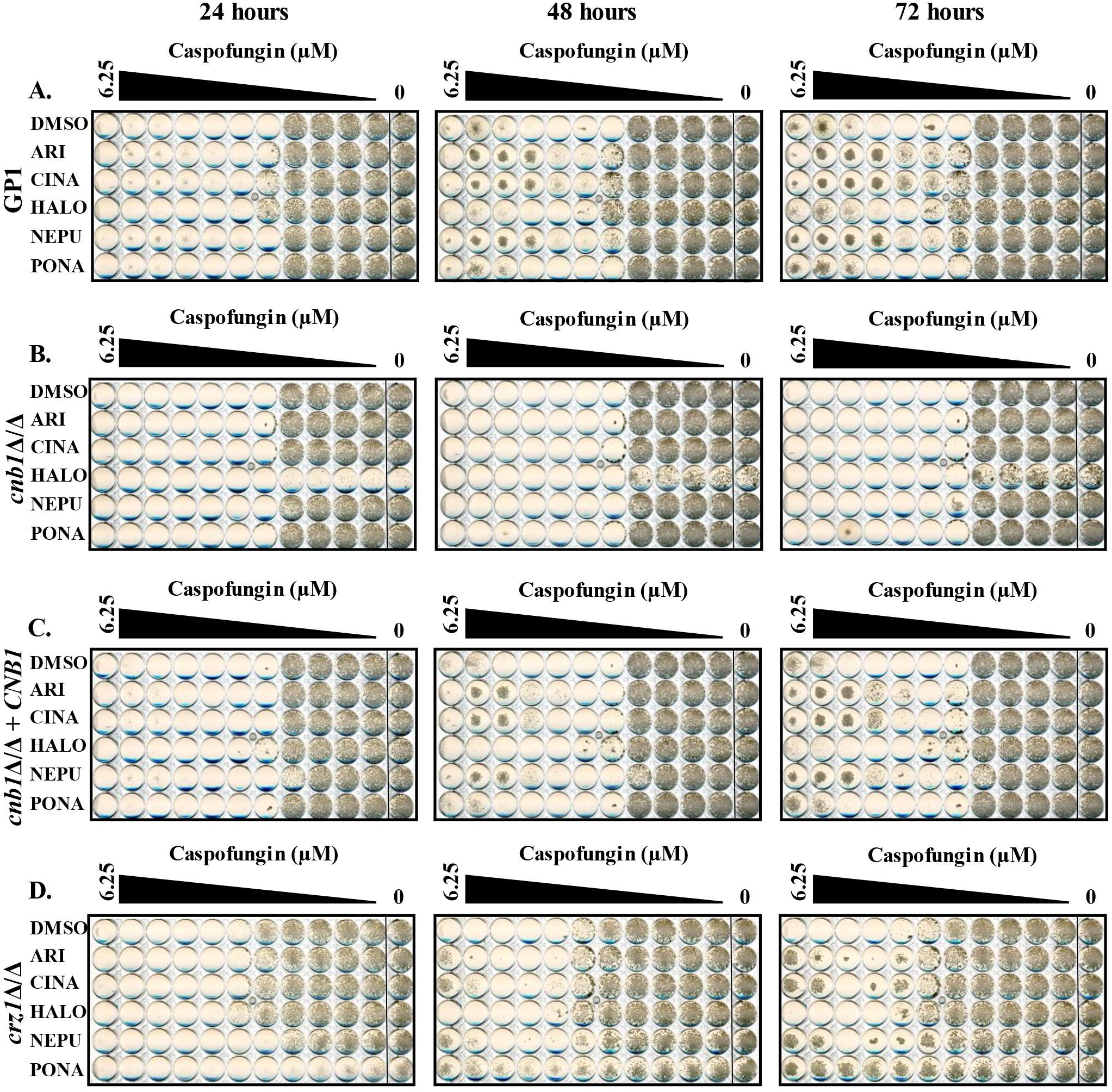
Drug echinocandin tolerance is dependent upon Cnblp. *C. albicans* wild-type strain GP1 **(A),** *cnblA/A* **(B),** *cnblA/A + CNB1* **(C),** and *crzlA/A* **(D)** mutants were grown with in RPMI-pH 7 (2% glucose) supplemented with either 5 μM aripiprazole (ARI), 5 uM cinacalcet (CINA), 5 μM haloperidol (HALO), 5 μM netupitant (NEPU), 2.5 μM ponatinib (PONA), or vehicle (DMSO) in combination with increasing caspofungin concentrations. Plates were incubated at 35°C, and imaged after 24, 48, and 72 hours. Images are representative of assays performed in biological duplicate.

**Figure S6.**
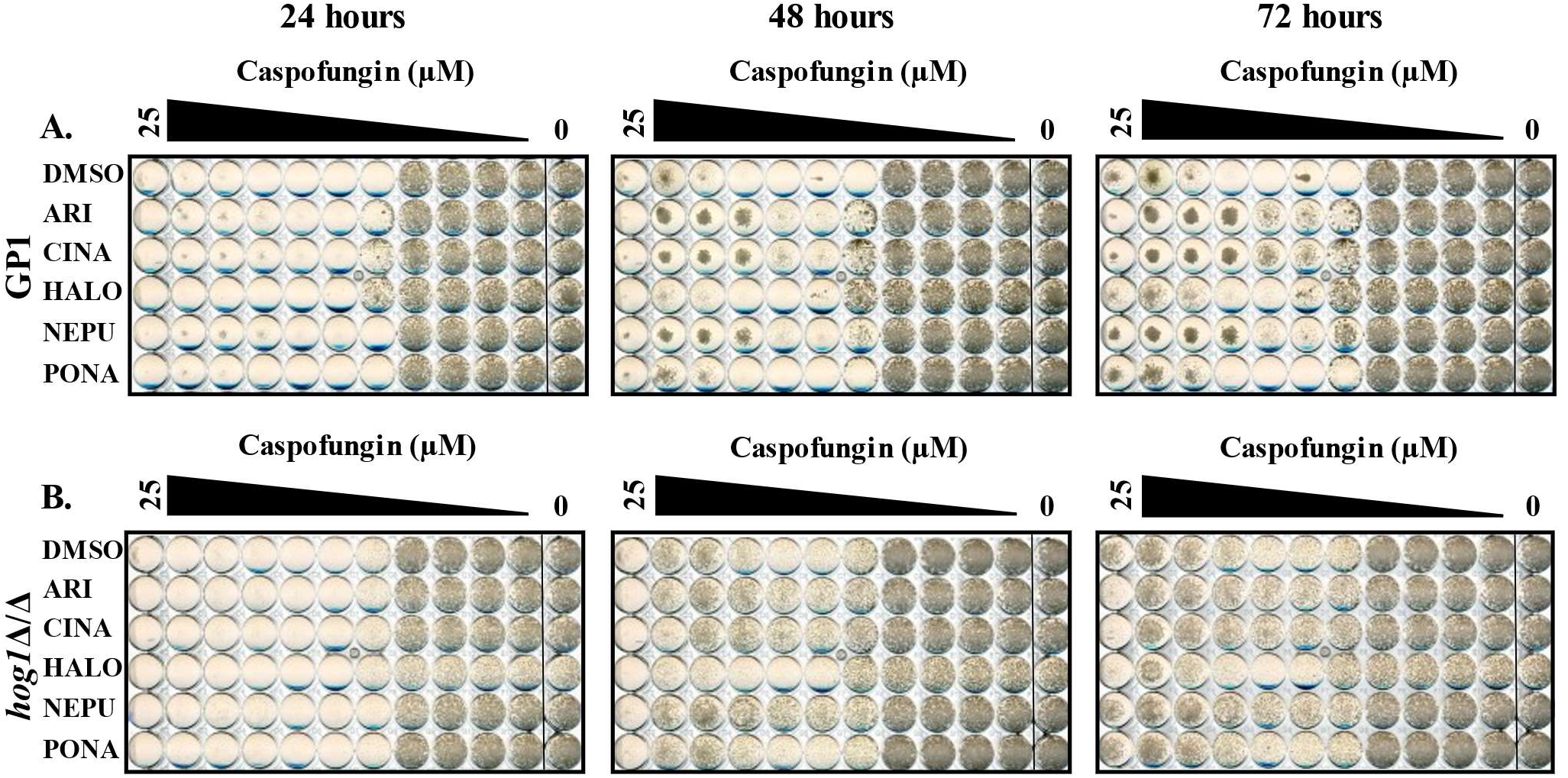
Deletion of *HOG1* induces echinocandin tolerance that is enhanced by antagonist treatment. *C. albicans* wild-type GP1 **(A),** and a *hoglA/A* mutant **(B)** were grown in RPMI-pH 7 (2% glucose) with supplemented with either 5 μM aripiprazole (ARI), 5 μM cinacalcet (CINA), 5 μM haloperidol (HALO), 5 μM netupitant (NEPU), 2.5 μM ponatinib (PONA), or vehicle (DMSO) in combination with increasing caspofungin concentrations. Plates were incubated at 35°C, and imaged after 24, 48, and 72 hours. Images are representative of assays performed in biological duplicate.

**Figure S7.**
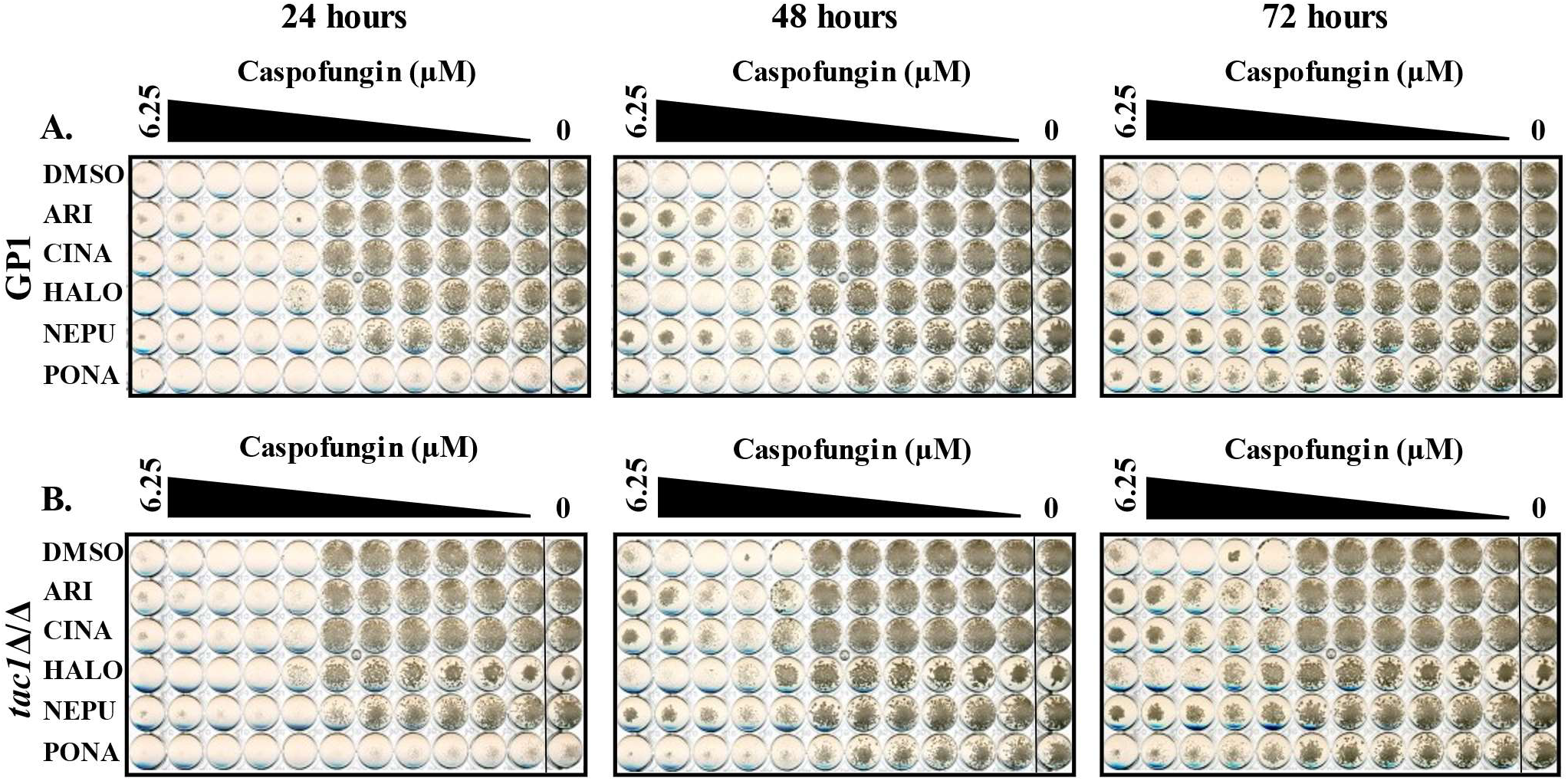
Echinocandhi antagonists are not dependent upon azole resistance mechanisms. *C. albicans* wild-type GP1 **(A),** and a *taclA/A* mutant **(B)** were grown in RPMI-pH 7 (2% glucose) with supplemented with either 5 μM aripiprazole (ARI), 5 μM cinacalcet (CINA), 5 μM haloperidol (HALO), 5 μM netupitant (NEPU), 2.5 μM ponatinib (PONA), or vehicle (DMSO) in combination with increasing caspofungin concentrations. Plates were incubated at 35°C, and imaged after 24, 48, and 72 hours. Images are representative of assays performed in biological duplicate.

**Table SI.**
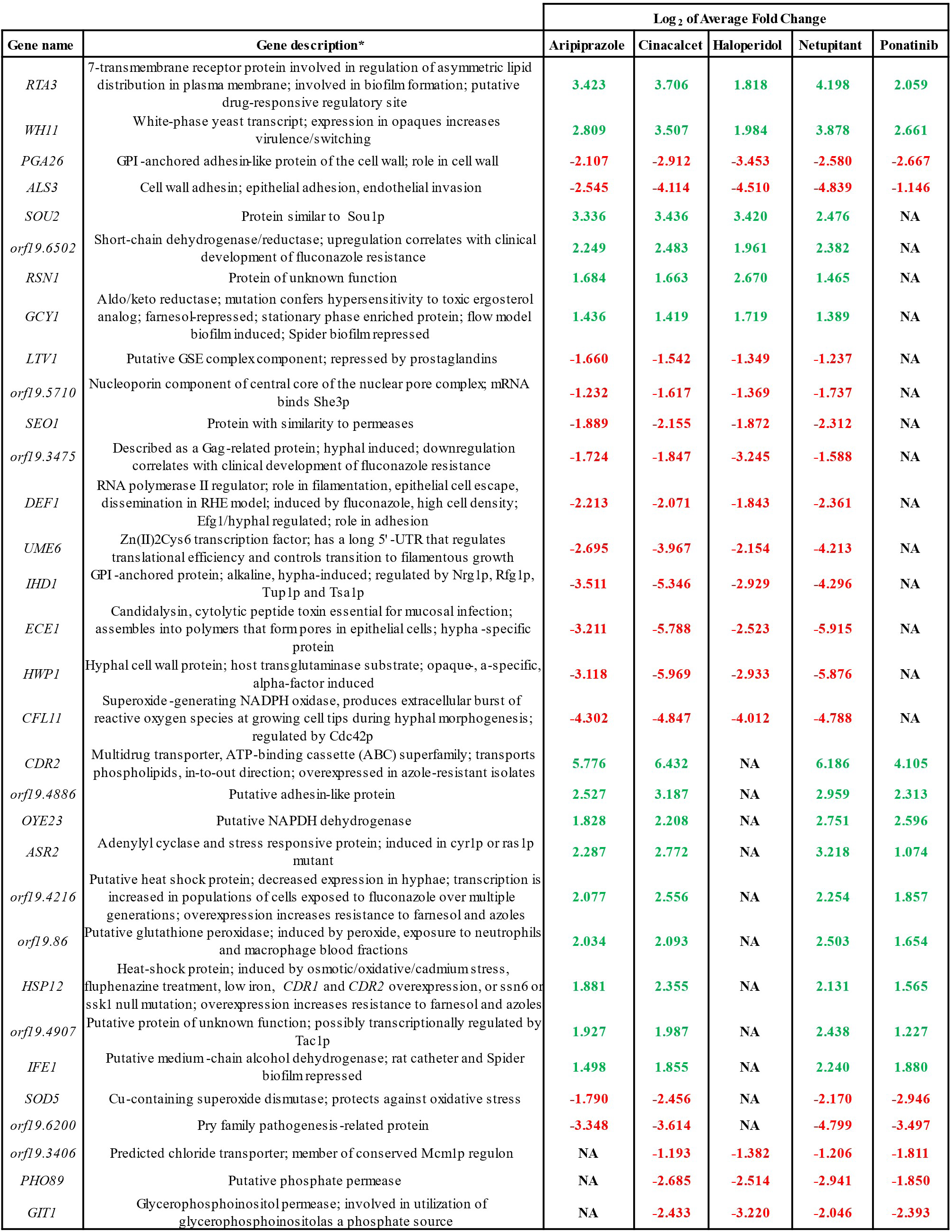
List of *C. albicans* gene transcripts responsive to echinocandin antagonists. *as described in Candida Genome Database. *non applicable (NA) where there was no significant change in transcript regulation compared to vehicle heated cells alone.

**Table S2.**
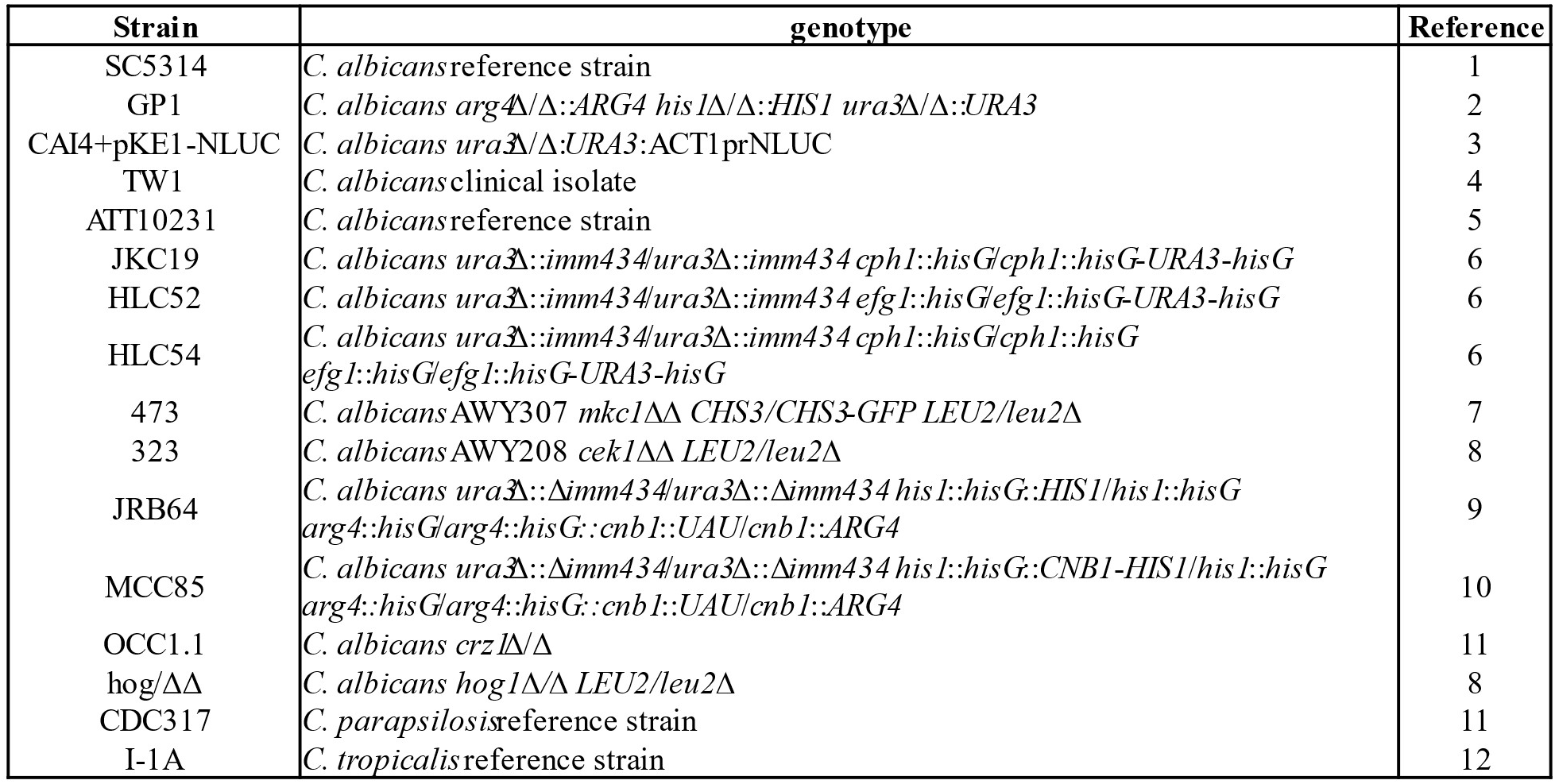
List of strains used in study.

